# Principal Genes: A PCA-based approach to highly variable genes selection for scRNA-Seq analysis

**DOI:** 10.64898/2026.08.07.743504

**Authors:** Enos Kakwambi, Tuan Nguyen, Suryaveer Kapoor, Moussa R Marmar

## Abstract

Single cell RNA-sequencing (scRNA-Seq) data are typically represented as cell-by-gene count matrices, which capture the expression of each gene as detected in the sampled cells; often a heterogeneous population of multiple different cell types or cell states. Almost all scRNA-Seq analysis workflows have a gene selection step prior to applying clustering algorithms which helps remove genes with low variability and hence reduce the high-dimensional gene space. A de-facto method for achieving selection of highly variable genes (HVG) uses dispersion and mean expression scores to evaluate the variability of each individual gene. However, methods based on direct mean-to-variance relationship for gene selection often suffer from susceptibility to variance instability and arbitrary determination of the optimal number of genes to use in downstream analysis tasks, additionally, they often prioritize genes with low abundance but high variance. Here, we propose an innovative method for selecting highly variable genes that is not based on mean to variance ratios: “Principal Genes (PG)” method; it utilizes the rotations (or loadings) from Principal Component Analysis (PCA) to calculate a novel variability score per gene that we name “Gene Principal Score (GPS)”. GPS helps evaluate the genes based on their contribution in the PCA rotations and hence ranks the genes according to their variability from highest to lowest variable genes. For efficient implementation we utilize Augmented Implicitly Restarted Lanczos Bidiagonalization methods to efficiently obtain Principal Components (PCs) associated with the largest variance. Genes with the highest GPS score, i.e. Principal Genes, can then be used for downstream analysis tasks, especially the clustering step. To test the performance of our highly variable gene identification method, we use several validation strategies, including clustering of labeled single cell RNA-Seq data (i.e. data with known ‘ground truth’ cell type labels). Furthermore, we measure the performance of our method against dispersion-based highly variable gene (HVG) selection approaches. We use several validation metrics, including sensitivity and adjusted rand index scores for clustering based on genes selected using our method against genes selected using HVG; and our validation datasets include six real labeled single cell RNA-Seq datasets. Our findings show that our new method, Principal Genes, is comparable and often favorable in performance in selecting highly variable genes and achieves ultra-fast gene selection from PCA results.

## 1 Introduction

Single-cell RNA sequencing (scRNA-seq) has become a leading technology used to analyze gene expression from thousands of genes at the level of individual cells in a massively parallel approach. In recent years, it has become one of the most used assays that allows researchers to study the heterogeneity and complexity of gene expression within cells’ populations, revealing unique cell types and states that might be missed by traditional bulk RNA sequencing methods. ScRNA-Seq datasets are typically represented as cell-by-gene count matrices, which capture the expression levels of each gene as detected in an often heterogeneous population of multiple different cell types or cell states. Prior to performing downstream analysis tasks such as clustering algorithms, most scRNA-Seq analysis workflows recommend a feature selection step which selects genes with high variance and high mean, i.e. selects Highly Variable Genes (HVG), reducing the high-dimensional gene space and improving the speed and accuracy of clustering and other algorithms in downstream analyses.

A de-facto approach for getting HVGs uses dispersion or mean expression versus variance scores to evaluate the variability of each individual gene and was first described in [12] for single cell RNA-Seq data. The approach is implemented in popular packages such as Seurat’s R package, which ranks and selects genes based on various mean-vs-variance methodologies [17].

Numerous studies explore the task of selecting highly variable genes from scRNA-seq data; however, similar to Seurat’s approach, existing methods almost exclusively focus on dispersion or direct mean-vs-variance approaches, giving only different flavors of the same basic recipe: mean expression and variance of genes are calculated and a formula utilizing both (e.g. with variance stabilization or binning, etc.) is calculated to score the genes. This achieves highly variable gene scoring based on highest dispersion or mean-vs-variance ratios or similar relationship of mean-vs-variance. Such approaches are represented and evaluated extensively in the literature such as in [22], [20], and [9]. Since our study is not a benchmarking study of methods based on dispersion or direct mean-vs-variance relationship, we refer the user to the cited literature for comprehensive benchmarking of these methods. Other scRNA-Seq pipeline evaluation and benchmarking studies that evaluated mean vs variance based gene selection methods and their effect on the overall analysis performance include [8] and [3], [21], [18] and other.

Another staple tool in scRNA-Seq data analysis workflows is Principal Component Analysis (PCA) which is a popular dimensionality reduction tool for high-dimensional data that projects data matrices onto new axes of greatest variance called Principal Components (PCs). It is typically used to examine variability and sometimes for clustering in lower dimensional space. The largest two or three PCs are used to present the data in a new two or three-dimensional space for easier visualization of the underlying structure of data [5]. PCA is a well studied approach; whether more classic PCA algorithms or tailored versions, specifically customized to suit the nature of scRNA-Seq data, various PCA benchmarking studies evaluate PCA techniques as an important part of the toolbox for single cell RNA-Seq analysis and point out the strengths and insights that can be obtained from PCA for single cell pipelines; the user is referred to [2], [13] and [18] for more on PCA approaches in scRNA-Seq.

We highlight that at the core of most PCA techniques, the PCs can be calculated via Singular Value Decomposition (SVD) techniques. Results from SVD return the magnitude of each gene’s contribution to variation captured via each PC. This is typically referred to as PCA’s *rotations* or *loadings* matrix.

Hence, we hypothesized that PCA rotations can be elegantly used to implement an alternative method to select highly variable genes that does not rely on a typical mean-vs-variance approach. We propose a novel method ***’Principal Genes (PG)’*** for highly variable gene selection that uses the variance information already captured in the PCA rotations to pick out the most variable genes. Instead of the dispersion-based analysis applied to the gene expression mean values, our method uses PCA rotation values to rank and sort genes based on their precise contribution to variability in the data.

In the following sections (2.1 to 2.4) we detail our algorithm for highly variable gene ranking and selection and in subsection 2.4.2 we introduce a new score for ranking Principal Genes. In section 2.5 we describe our validation datasets and methodology followed by our results and discussion in section 3 and we evaluate our methods time performance in section 3.5 and conclude in section 4.

## 2 Materials and Methods

### 2.1 Overview of Principal Genes Method

In PCA, rotations represent the correlation between the original variables, i.e. genes for scRNA-Seq, and the calculated principal components. They indicate how much each gene contributes to a particular principal component. Rotations or loadings are essentially the coefficients in the linear combination of original variables/genes that create the principal components. We hypothesized that extracting the participation of each gene in each principal component will be a strong indication of this genes’ variability property and can be used to rank and score genes for gene selection.

Our method starts by filtering out commonly known housekeeping genes that tend to be ubiquitously expressed across cells. This is done by removing genes that have zero variance in the data, followed by removing housekeeping genes by naming convention. Following QC and filtering, we perform PCA (which can be preceded by a step to determine the optimal number of PCs), followed by extracting each gene’s rotation or loading score in each PC where the gene is participating. This is further aggregated for each gene across all participating PCs into a combined new proposed score, Gene Principal Score (GPS) (described in detail in section 2.4.2) that quantifies the variability contribution of each gene. Genes are then ranked by the GPS score and genes with highest GPS are picked into the highly variable Principal Genes set. This set continues to grow, until a total percentage cutoff is met. The cutoff represents the desired fraction of the total contribution of variance that can be captured by the PCs. The selected Principal Genes set can then be used in further downstream analyses or other steps as customary in a typical scRNA-Seq workflow.

### 2.2 QC and Pre-processing

Primary filtering performed for all datasets and for both methods Principal Genes and Seurat approaches selects genes with zero total expression or zero variance for elimination. To reduce the size of the the human primary motor cortex data (detailed in 2.5), we filtered out genes detected in less than 10 cells for all methods. Each method is then used in their respective recommended workflow for fairness. For Principal Genes, we recommend filtering out housekeeping genes, mitochondrial and ribosomal genes as well as any unknown genes in a cleanup step that can be customized by the user. The filtering step or regressing out housekeeping or mitochondrial genes, etc. is a standard best practice of the QC workflow for single cell RNA-Seq data [12] [14]. For our experimental testing we report both results, with and without this filtering step, with filtering results are reported in 3 and without filtering reported in supplementary in 0.3.

### 2.3 Selecting an ‘Optimal’ Number of PCs

PG method utilizes the first *’few’* PCs capturing the highest variability. Selecting the ‘optimal’ number *k* of PCs to select is still an open problem, however, there are a number of recommended heuristic strategies. One demonstrated strategy is the use of the so-called ‘elbow’ plot (see Figure 1 for an example) of the standard deviation or singular value of each PC, where selecting a *k* number of components based on where the plot starts to level off is a popular method for selecting an optimal number of PCs.

**Figure 1.**
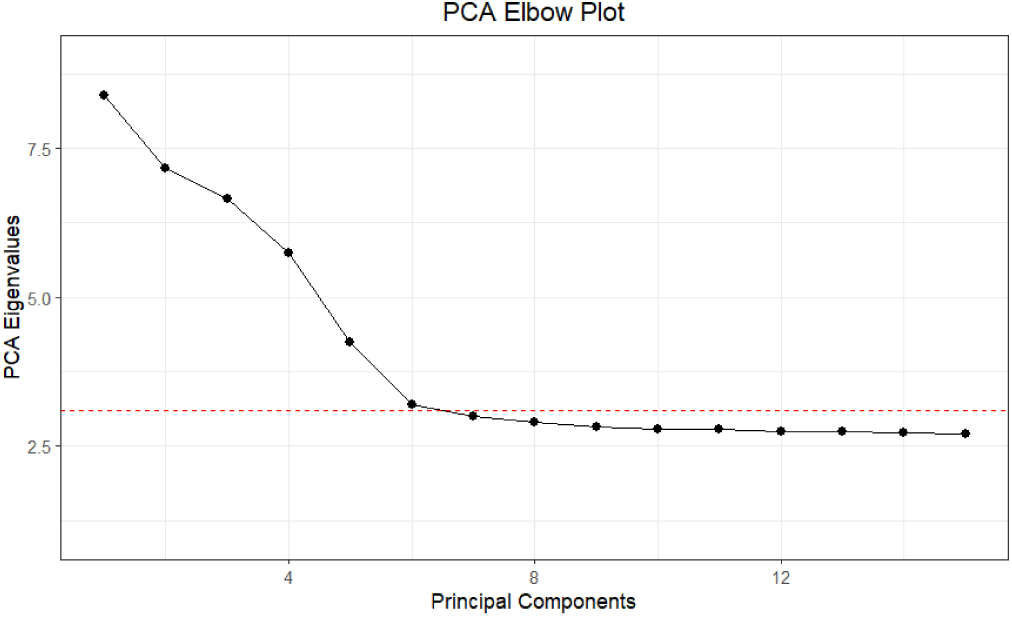
Elbow plot example, the PBMC dataset used in this plot is described further in section 2.5.) The horizontal line cuts the curve in or around the ‘optimal’ number of PCs to select.

One can notice that after this proposed cutoff at the ‘elbow’ point, the singular values get closer and closer together suggesting diminishing returns. This approach usually helps to visually determine a suitable cutoff, but here, we employ a quick elbow algorithm based on [7] that computes the optimal number of PCs by first converting PC eigenvalues as proportions of total variance, followed by identifying components that contribute to a minimum (heuristically determined) proportion of variance and finds the largest relative drop in adjacent PC variance proportion, electing the consecutive principal component’s index as the value for *k*. This ensures a near optimal selection of the number of PCs without requiring manual or user input.

### 2.4 Principal Genes Selection

#### 2.4.1 Calculating PCs and Rotations

A standard SVD algorithm can be used to generate the PCs and needed rotations’ matrix, however, to achieve superior performance, we use the Implicitly Restarted Lanczos Bidiagonalization (IRLBA) algorithm [4] as a method for estimating the largest principal components in the data.

This algorithm utilizes the fact that the first few PCs typically have the largest singular values and therefore is focused on computing a few of the largest singular values and corresponding singular vectors of a large matrix. It quickly and accurately estimates the largest singular values of the scaled and centered a gene by cell matrix, *A ∈* ℝ*^g×c^*, where *g* = #*ofgenes* & *c* = #*ofcells*. The transposed counts matrix *A^T^* is passed to the algorithm with *k* selected principal components for the standard decomposition in PCA given by:

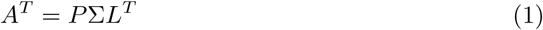

- **P** *∈* ℝ*^c×k^* with *k* principal components and *c* embeddings representing *k* coordinates for each cell
- **Σ** *∈* ℝ*^k×g^* with the diagonal being ordered eigenvalues for each *k* principal component where *σ*_1_ *> σ*_2_ *> . . . > σ_k_*, representing the variance contribution of each principal component
- **L^T^** *∈* ℝ*^g×k^* with rotations/loadings representing the contribution of gene *g* to principal component *k*

We then utilize the matrix **L^T^** to extract a measure of gene ‘importance’ as described in 2.4.2.

#### 2.4.2 Gene Principal Score

To select and prioritize genes based on their contribution to implicit variance contained within the calculated components, we created a new metric, Gene Principal Score **(GPS)**, which we define in Equation 2:

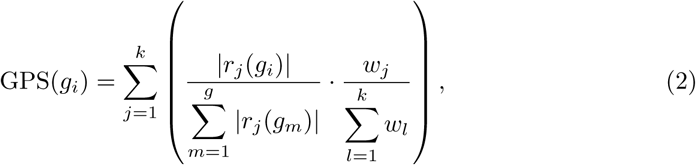

Where:

- *r_j_* is the rotation/loading vector of PC *j* and represents the *j − th* column in **L^T^** -note that the gene’s variance contribution can be in either direction of the PC but only the magnitude of the contribution is relevant to the score.
- and *w* is the diagonal vector of Σ

This definition ensures that every gene’s rotation value in each Principal Component is scaled for that particular PC before factoring the proportion of variance which that PC explains. These scaled intermediate scores are then summed over all PCs per gene to compose the final GPS value per gene. The GPS scores of the genes are sorted in descending order in the last step of the algorithm where genes with higher GPS scores are ranked higher for selection.

### 2.5 Validation Approach

#### Datasets

To test and validate our new approach, we applied the Principal Genes method to six real and publicly available scRNA-Seq data with known labels (i.e. ground truth cell type labels). First, the labeled and FACS sorted Peripheral Blood Mononuclear Cell (PBMCs) dataset from 10x Genomics [1] platform, obtained from [14], we also sample the FACS sorted Lung and Liver datasets from Tabula Muris Consortium [6] creating two additional labeled sets. We use a dataset obtained from [15] where Pollen et. al manually curated labels representing the ground truth for a marker-gene–based manual annotation of a developing human cerebral cortex. Additionally, we subset the Allen Institute’s data on the human primary motor cortex (HPMC), selecting five labels from the original dataset. Labels were assigned through unsupervised transcriptomic clustering of the gene expression data [10]. Finally, we use a sampled mouse motor cortex single-nucleus data from Azimuth Hub Consortium (Azimuth MMC), labels for this set were obtained via marker-gene expression analysis [19]. The MMC dataset (11 cell types in true cluster labels) was further used in section 3 as a case study to show the importance of selected genes in other downstream tasks such as heatmap visualization and enrichment analysis. Figure 2 shows the PCA plots of the datasets used in our study.

**Figure 2.**
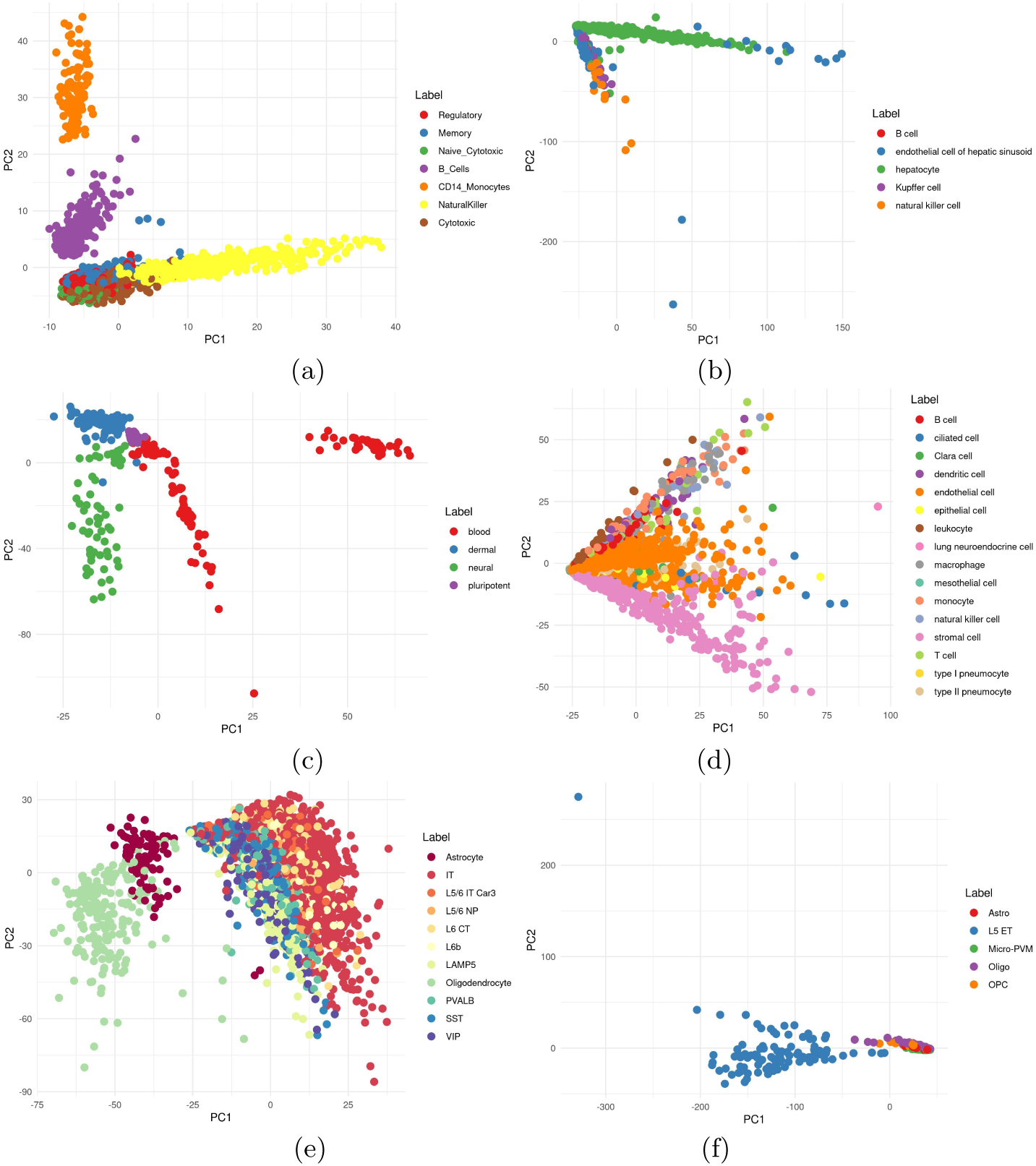
a) PCA plot of a) PBMCs, b) Tabula Muris Liver, c) Pollen, d) Tabula Muris Lung, e) Azimuth MMC and (f) Human Primary Motor Cortex using PC1 and PC2. Ground truth cell types from [14], [6], [15], [6], [19], [10] respectively are used for color coding.

#### Clustering Accuracy

Clustering for cell type identification is an essential step in scRNA-Seq analysis workflows that occurs downstream from highly variable gene selection. Therefore, to test and validate our new selection method, we used spherical-k-means clustering and evaluated the overall accuracy metric, i.e. sensitivity or total number of True Positives -by majority assignment- divided by the Total number of True Cell Labels. This approach is highly interpretable and is used to determine how well the algorithm performed in selecting highly variable genes that can be efficiently used as informative genes in downstream analysis tasks.

It is important to note here that clearly other clustering algorithms can be used for this evaluation as long as the same clustering approach is used for all comparisons. We compare with Seurat’s HVG selection as representative of the mean-vs-variance methods in section 3 and further with Seurat’s HVG with variance stabilization in 0.2.

Furthermore, we use adjusted rand index (ARI) [16] as a measurement of pairwise agreement between cells in the unsupervised clustering results compared to the true labels. ARI gives a maximum score of 1 where 1 shows perfect agreement between the unsupervised clusters and the true labels, 0 on the other hand shows an agreement level expected by random clustering and negative values shows less agreement than expected by random cluster assignment.

#### Optimal number of genes to select

The proposed Principal Genes algorithm can assign a GPS score value to every gene in the dataset and a list of genes sorted in descending order based on their GPS scores is returned by the proposed Principal Gene algorithm. This ranked list is further used to select the ‘top’ or highly variable genes from the list of all given genes.

Instead of using an arbitrary cutoff on the number of ‘top’ genes selected, we define the top genes - Principal Genes - as all genes with cumulative GPS score *<*= *c*, where *c* is a cutoff value that can be set by the user, representing the desired total cumulative variance contribution that is to be covered by the selected genes. As an example, a cutoff of 20% translates to selecting all genes whose contribution amounts to 20% of implicit variance contribution across all PCs. This approach provides an interpretable parameter for the method and does not force the user to ‘guess’ an optimal number of genes to be selected but provides the final list of Principal Genes to match the selected desired cumulative variability contribution. Note, to study the effect of setting this cutoff parameter, we vary the cutoff based on cumulative GPS scores in the results section 3 from 10%-50% with increments of 2% and select the top *x* number of genes needed to achieve the desired cutoff. Each group of *x* genes corresponding to each cutoff value is used as the top highly variable genes for clustering analysis and the accuracy scores are noted (see next section 3 for results details).

## 3 Results and Discussion

### 3.1 Accuracy Results in Downstream Analysis

#### Clustering Sensitivity

Clustering is a staple task in single cell RNA-Seq downstream analysis workflows. To evaluate the precision of clustering when selecting the top highly variable genes using the Principal Genes method, we first used the PBMCs dataset and obtained the optimal number of PCs (= 6 using the elbow algorithm as described previously 2.3). Using the top 6 PCs we obtained the ranked list of Principal genes for any given cumulative cutoff value and calculated the overall accuracy score for the 7 true cluster labels as well as the ARI score. We performed the same analysis for all datasets from Figure 2 a to f. Varying the genes selected for use in sk-means clustering based on their cumulative score produces the set of overall accuracy/sensitivity values seen in Figures 3 and the ARI scores seen in 4) for the PBMCs, Liver, Pollen, Lung, MMC and HPMC datasets respectively.

**Figure 3.**
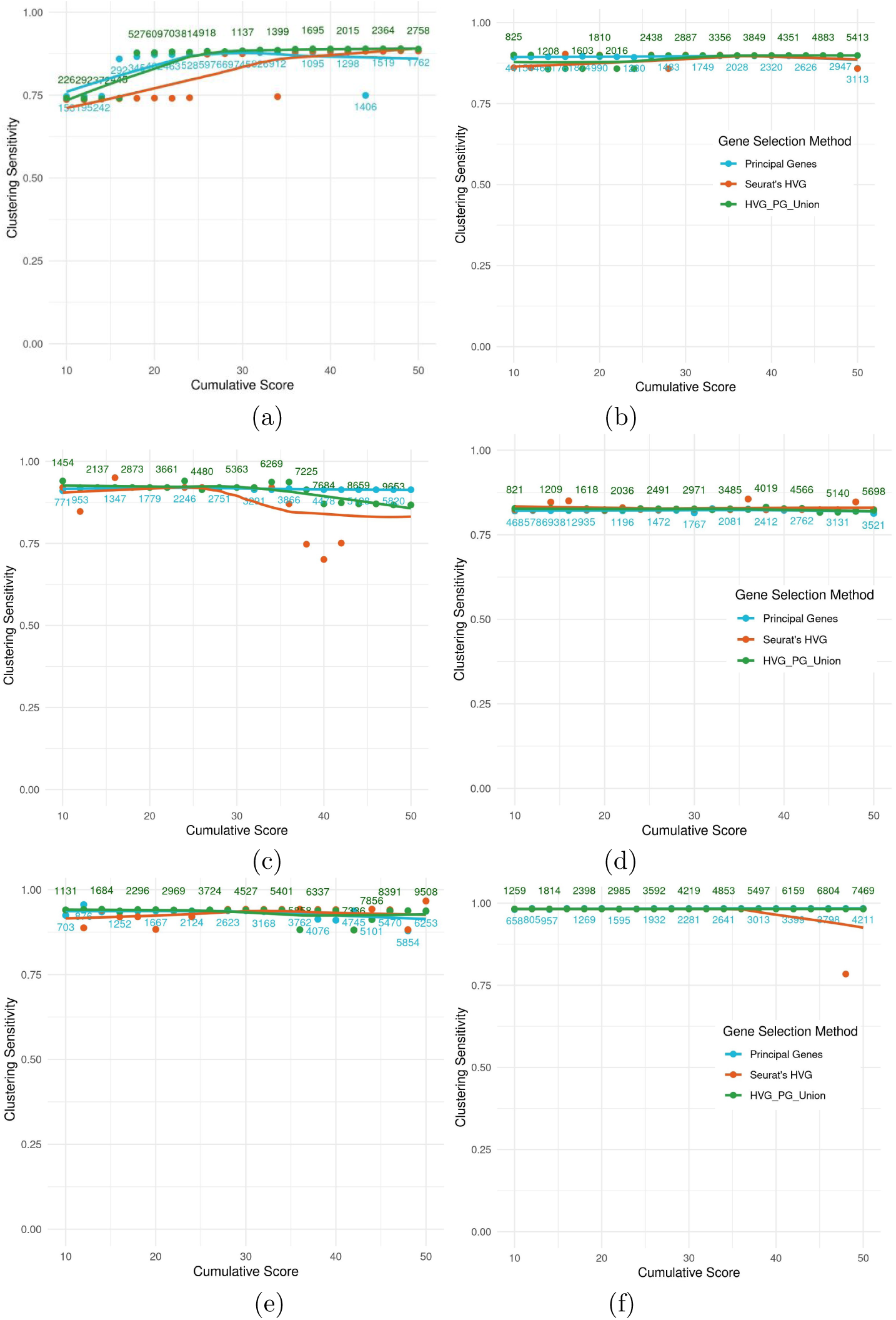
SK-Means Clustering Sensitivity comparison between Principal Genes, Highly Variable Genes, and the union of both for a) PBMCs, b) Tabula Muris Liver, c) Pollen, d) Tabula Muris Lung, e) Azimuth MMC and (f) Human Primary Motor Cortex. PGs selected for clustering are ordered in descending order and subset into groups of increasing cumulative GPS. Top Genes from Seurat’s HVG are selected to match the number of genes in each group. The union combines unique genes selected by both methods at each point. The number of genes at each group are shown in blue for HVGs and PGs and in green for the union.

Peak sensitivity score of (0.89) for the PBMC dataset can be observed; selecting a cutoff of only 20% cumulative score or higher already achieves near peak accuracies with only 402 Principal Genes selected to achieve this high sensitivity score. All other datasets follow a similar trend with peaks in their sensitivity scores appearing with smaller cumulative scores. It is important to note, that for the other datasets, peaks are reached much ‘earlier’ in their cumulative score. This can be attributed to the higher number of genes present in the subsets compared to PBMCs. To compare with the dispersion-based ‘traditional’ HVG approach [12] [17], we calculated the overall accuracy/sensitivity scores using an equivalent number of HVGs - obtained by Seurat’s R Package implementation of HVGs [17]. In all datasets, we observe higher or similar sensitivity scores for Principal Genes method and when using the union of both methods.

#### Adjusted Rand Index

Furthermore, ARI scores convey similar results, with all datasets showing ARI results from Principal Genes similar or higher than those of Seurat’s HVG, except for one dataset, the Azimuth’s MMC data where HVGs ARI scores are higher than PG but only when higher number of genes are selected. Nevertheless, Principal Genes method still outperforms in the clustering sensitivity score for the same dataset. Slight variability is observed in ARI measure compared to sensitivity scores. Overall, the performance of genes selected by Principal Genes is still similar or favorable to HVG scores in all datasets in at least one of the two measures if not both.

#### Bootstrapping

To strengthen the statistical rigor of our results, we performed bootstrapping of the data, sampling 80% of each dataset over five separate iterations followed by performing the clustering and metric evaluation steps. Bootstrapping ensures any stochasticity is accounted for in the resulting confidence intervals. Figure S1 shows that observed sensitivity scores results are indeed consistent across bootstrapping results of the replicated experiments; in fact for datasets such as PBMCs, Liver, Pollen, and MMC, Principal Genes show more stability in results represented by narrower error margins than observed in Seurat’s results.

#### Variance Stabilization

Variance stabilization transforms are sometimes applied in single cell analyzes to ‘stabilize’ variance across genes and leave the comparison focused on mean expression values [17] [11]. We examined the performance of Principal Genes against that of Seurat’s stabilized variance modality for highly variable gene selection for the Liver data. Results shown in S2 and S3 are consistent with better performance for Principal Genes; surprisingly, highly variable gene selection using Seurat’s stabilized variance approach did not result in better accuracy scores against Seurat’s own dispersion based approach. This can be attributed to loosing genes due to their diluted variance effect resulting from the stabilization transform.

### 3.2 Effect of Number of PCs Selected

In a previous section (2.3), we presented how the elbow method can be effectively used to set an ‘optimal’ number of PCs, however this algorithm requires calculating more PCs than needed to discover the elbow point, hence can be computationally expensive when computing resources are limited. For this reason and since the selected number of PCs is a configurable parameter in calculating the first step of our algorithm (IRLBA-based matrix), we examine the effect of selecting different numbers of PCs as input for our approach. In Figure 5 a) for PBMC dataset and b) for MMC we vary the number of PCs used and calculate the sensitivity scores for both sets respectively. Limited variability in the accuracy results can be observed, both sets show relatively stable accuracy values between 0.75 and 0.9 for PBMCs and between 0.8 and 0.965 for Azimuth. In general, we can see that there is no perceived advantage in terms of accuracy when a high number of PCs is selected since comparable accuracy scores are achieved with lower than 10 PCs.

**Figure 4.**
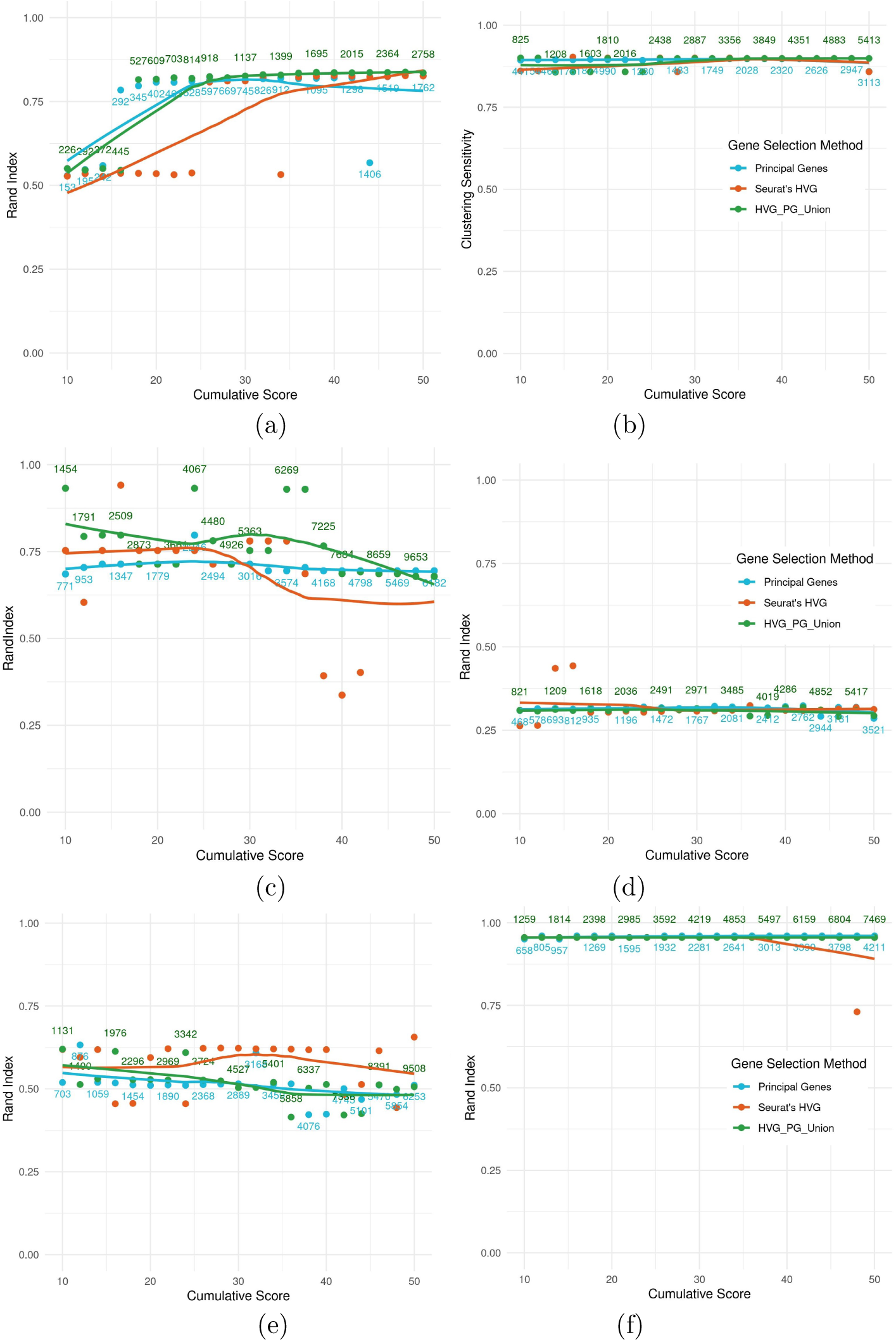
SK-Means Clustering ARI comparison between PG, HVGs and the union of both for a) PBMCs, b) Tabula Muris Liver, c) Pollen, d) Tabula Muris Lung, e) Azimuth MMC and (f) Human Primary Motor Cortex. PGs selected for clustering are ordered in descending order and subset into groups of increasing cumulative GPS. Top Genes from Seurat’s HVG are selected to match the number of genes in each group. The union combines unique genes selected by both methods at each point. The number of genes at each group are shown in blue for HVGs and PGs and in green for the union.

**Figure 5.**
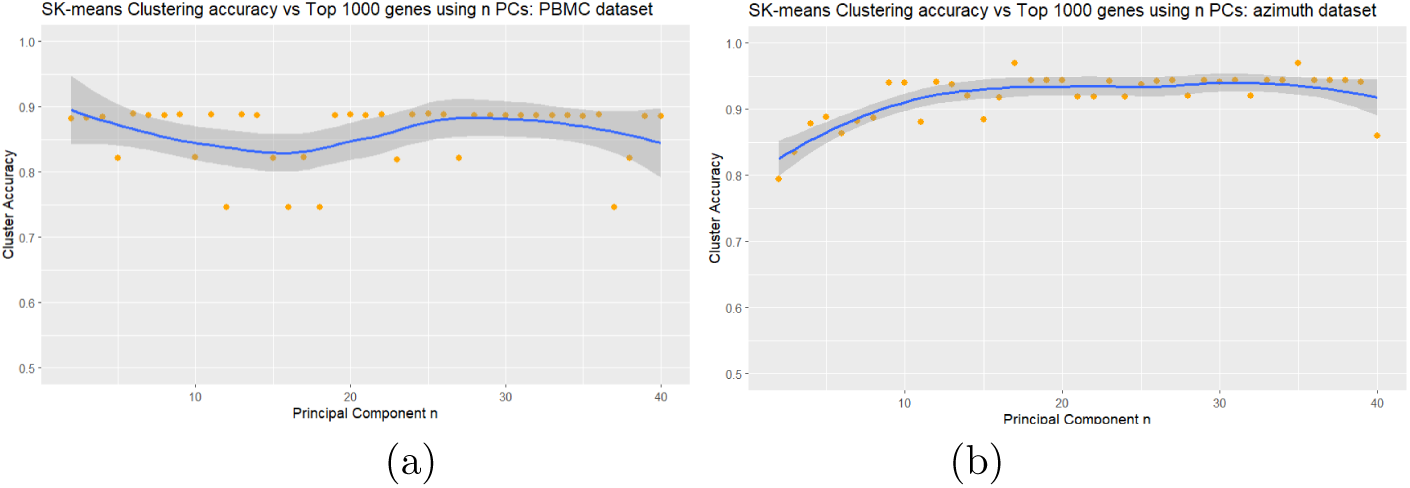
a) SK-means Clustering Sensitivity for PBMC’s with varying number of PCs chosen to calculate GPS scores (top 1000 Principal Genes were selected). b) Clustering Sensitivity for Azimuth’s Mouse Motor Cortex with varying number of PCs chosen to calculate GPS scores (top 1000 Principal Genes were selected).

### 3.3 Testing Distortion-Free PC selection

Based on the findings of [13], we wanted to examine whether or not Principal Genes might benefit from an implementation of distortion free principal component selection. Testing on various datasets revealed that distortion-free selection focused on finding principal components with z-normalized loading scores that follow a normal distribution, discarding ones that do not. Figure 6 panel (a) shows that the data does not follow the expected normal distribution. Important genes (the top 9 shown in Figure6 panel (b)) that are ranked highly within these non-normal principal components would potentially be discarded under the distortion-free scheme. These results suggest distortion-free PC selection is not advantageous in Principal Genes selection.

**Figure 6.**
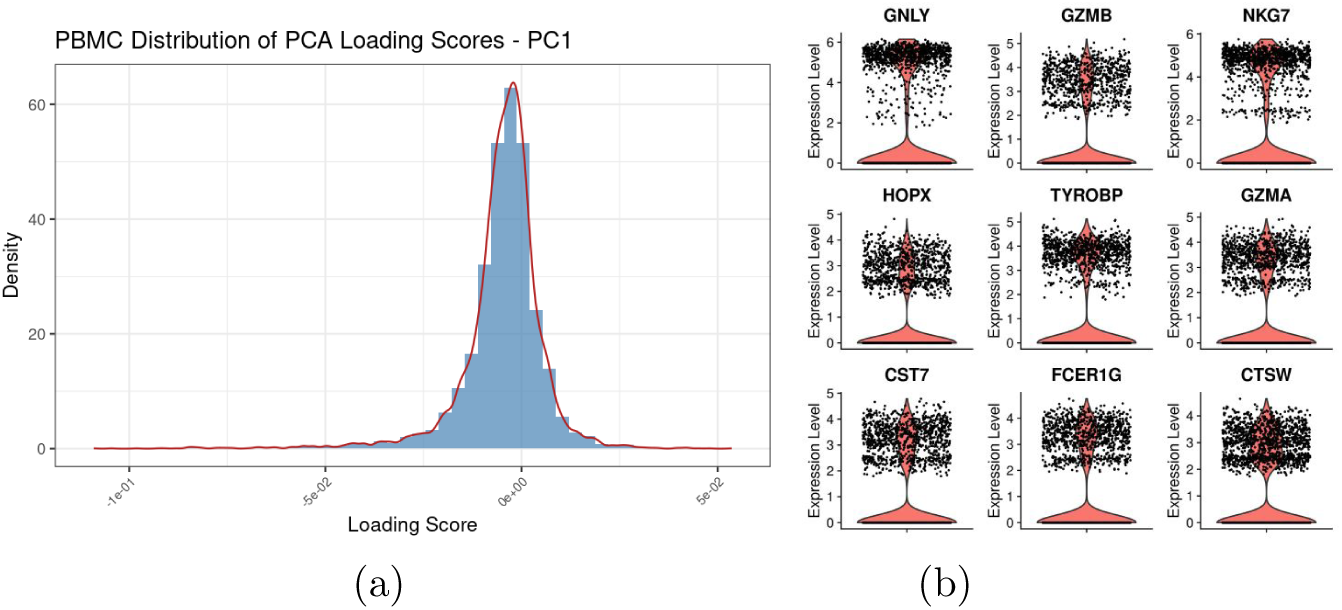
Histogram of the first principal component’s z-normalized loading scores obtained via IRLBA for a) PBMCs, and expression level violin plots of genes with the highest scores from PC1 for the b) PBMC dataset.

### 3.4 Principal Genes Properties Analysis

#### Top Gene Selection

To study the properties of our selected genes, we first ask whether a large intersection set exists between top genes based on GPS score,i.e. Principal Genes, versus when top variable genes selected based on dispersion analysis for HVGs. In Table 1 and 2 we show the top 25 Principal Genes (ordered list) and their ranks from dispersion-based HVGs selection for PBMCs and MMC datasets respectively; 57 genes out of the top 100 were selected by both methods for PBMCs dataset and 23 genes were shared for the MMC data. Furthermore, we observe that genes when ordered by variance in the tables S6 and S7, were ranked overall higher by Principal Genes’ GPS score than by HVG for both datasets.

**Table 1.**
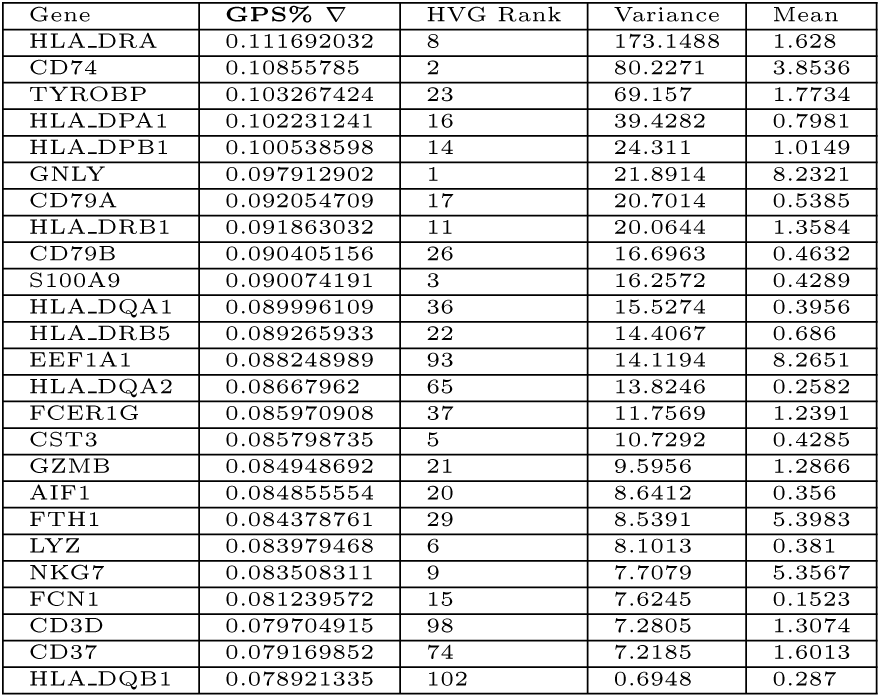
Top 25 PBMC Principal Genes vs rank in top 100 HVGs + Mean and Variance.

Interestingly, we note that several genes (e.g. HLA-DQB1 from PBMCs, CHN1 and CELF2 from MMC) selected by Principal Genes and ranked in the top 25 Principal Genes with clearly high variance and high expression mean scores were not picked up by dispersion-based HVG method and furthermore did not rank in the top 100 of HVG gene list.

#### Exclusive Gene Selection

To further examine the properties of the selected genes from each method, we note that other than shared genes, several genes are indeed exclusive to each method; the top genes selected exclusively by either method as well as the top genes selected by both as shared set are illustrated in Figure 7 in a GPS vs Dispersion plot and further via violin plots in Figure 8 (Violin Plots for other datasets are shown in Supplementary Figures S9 to S10).

**Figure 7.**
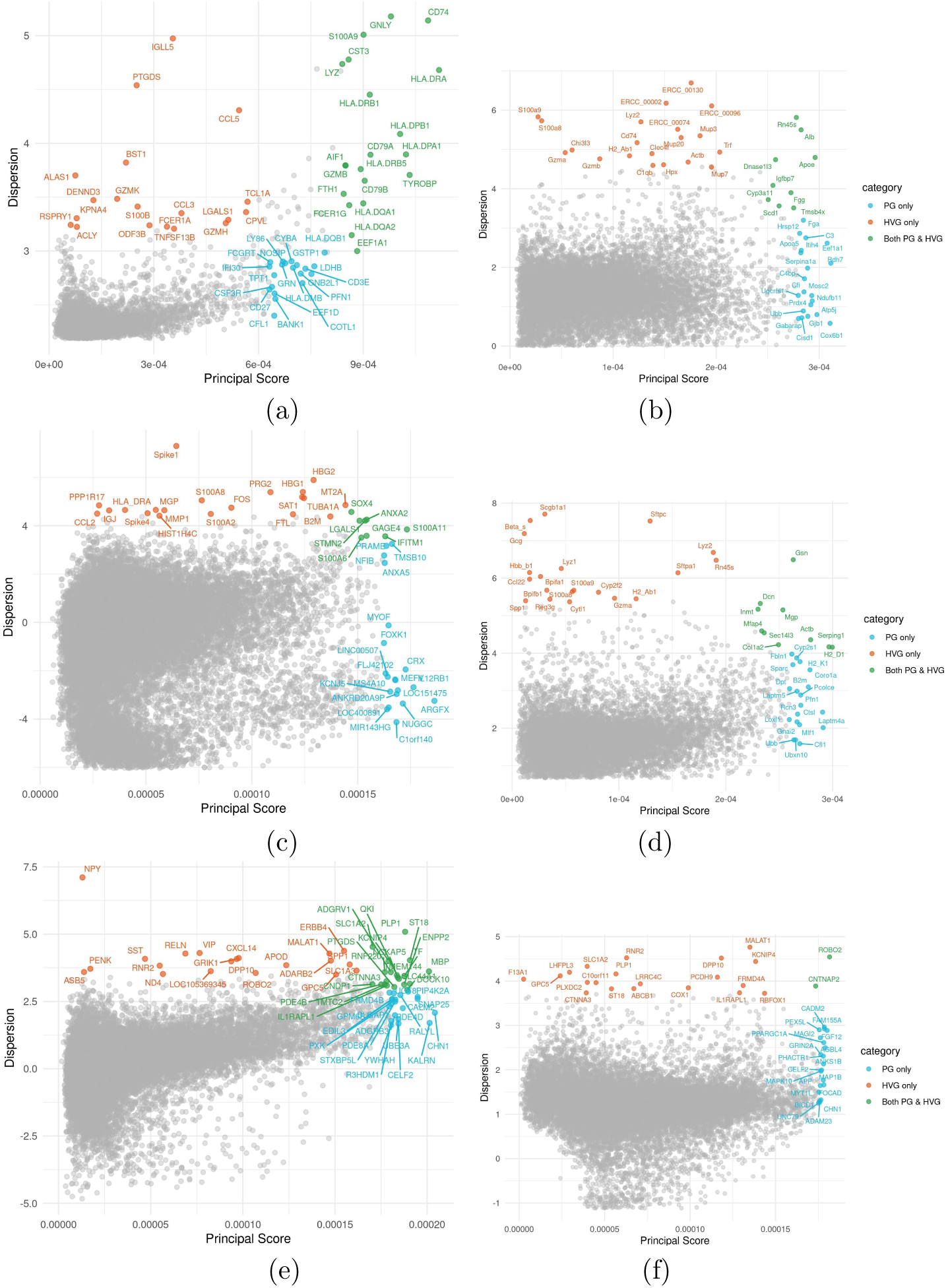
Gene Principal Score vs Dispersion — Top 20 genes highlighted by method (blue for PG exclusive, orange for HVG exclusive and green for shared gene selections for: a) PBMC dataset, b) Tabula Muris Liver dataset, c) Pollen, d) Tabula Muris Lung, e) Azimuth MMC and (f) Human Primary Motor Cortex.

**Figure 8.**
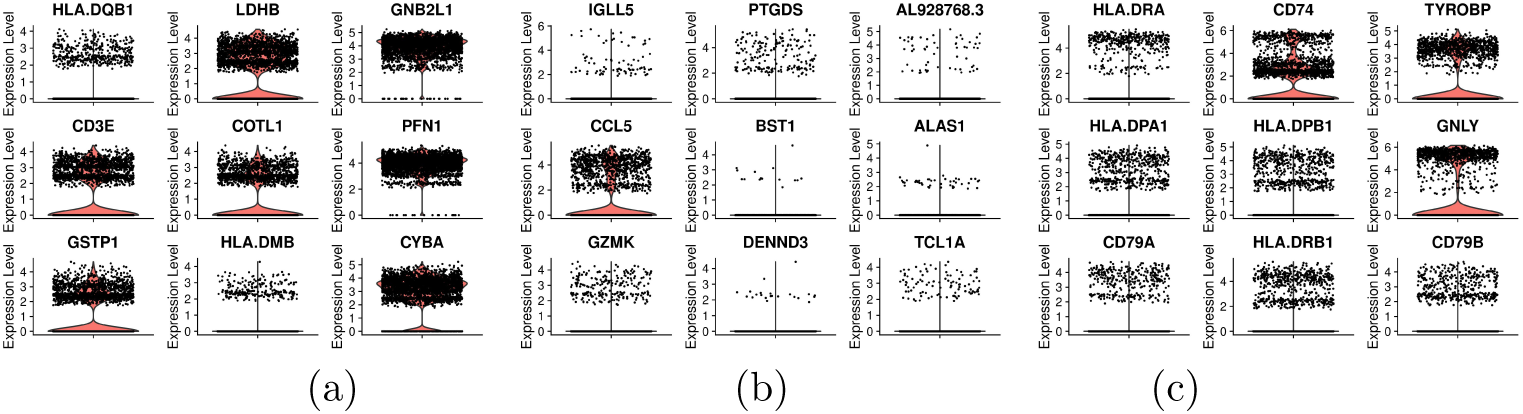
PBMC Dataset: a) Violin Plots for top 9 genes in PG’s top 100 absent in HVG’s top 100 (exclusive to PG in top 100). b) Violin Plots for top 9 genes in HVG’s top 100 absent in PG’s top 100 (exclusive to HVG in top 100). c) Violin Plots for 9 genes in the intersection of HVG’s and PG’s top 100 (shared within the top 100).

#### Expression and Variance Analysis

Interestingly, in general when comparing Principal Genes versus dispersion-based HVGs, we observe comparable clustering accuracy results, despite both methods prioritizing different gene sets (the overlap observed in the top 100 selected genes was between 23% and 50% and increases as the number of selected genes increase). The difference in genes selected by the two approaches points to both methods targeting and capturing different aspects of variability. As noted before, Figure 7 shows where the respective top selections of each method lie in a dispersion vs GPS score plot and in section 3 the performance of the combined gene sets in clustering tasks show an enhanced performance when combining both approaches.

To further analyze the statistical differences between the top genes selected by each method, we examine the average expression abundance and variance of each selected set of genes. Figure 9 shows boxplots for the mean and variance of the log2p normalized expression of the top 100 genes selected by each method across all used datasets. This analysis shows that Principal Genes method overall selects genes with higher average expression levels than HVG (Figure 9 a). The variance values of the top genes follow the same trend of higher averages (Figure 9 b) which the exception of the Pollen dataset, where the mean variance of PG is lower than HVG.

**Figure 9.**
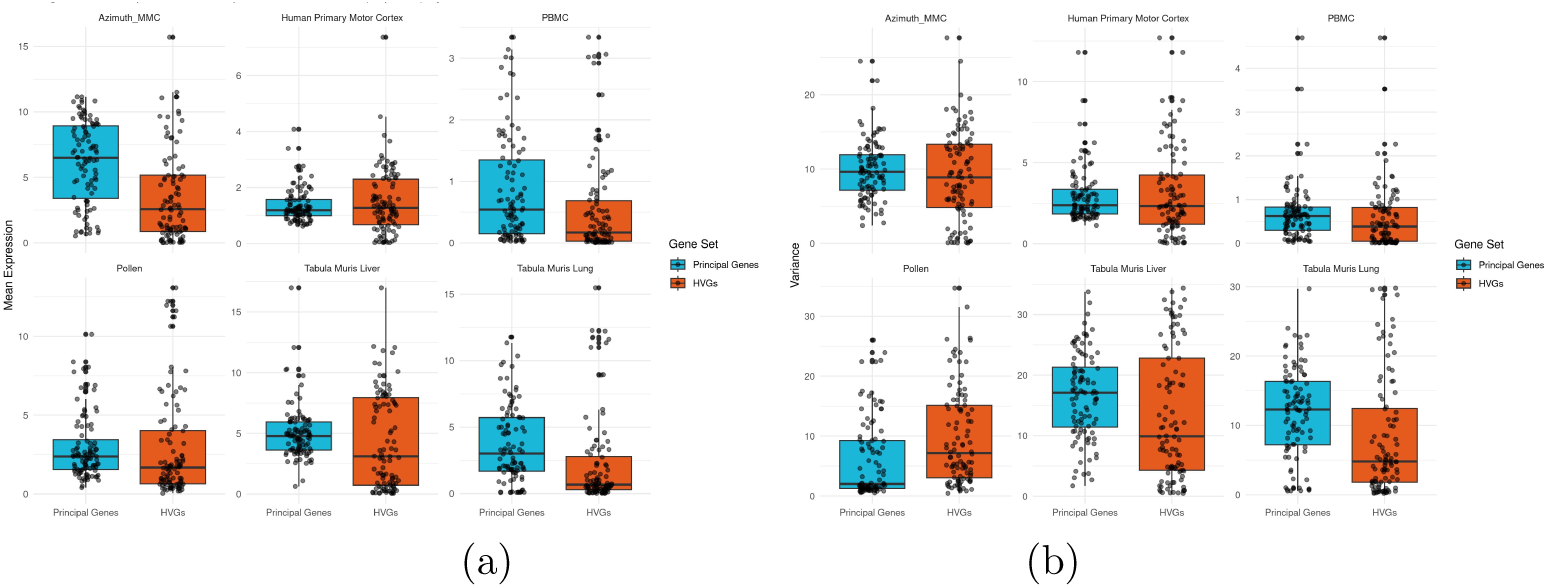
a) Boxplot of mean expression levels of top 100 selected Principal Genes and Highly Variable Genes across all datasets. b) Boxplot of variance of top 100 selected Principal Genes and Highly Variable Genes across all datasets.

#### Rank Correlation

To assess the correlation of ranks between the top selected genes by each method, we apply Kendall Rank Correlation Coefficient for the top 100 genes selected by each method across datasets (Table 3). All correlations were significant (p-value ¡ 0.05) except for Pollen dataset, and we find that for datasets with the smallest p-values, the *tau* values (correlation) ranges do not exceed 0.52 suggesting limited correlation exists between the gene selection approaches for the top ordered genes. This further confirms while both methods can capture genes that ultimately contribute to high performance in clustering and downstream analysis, both have their unique selection biases as described before in Figure 9.

**Table 2.**
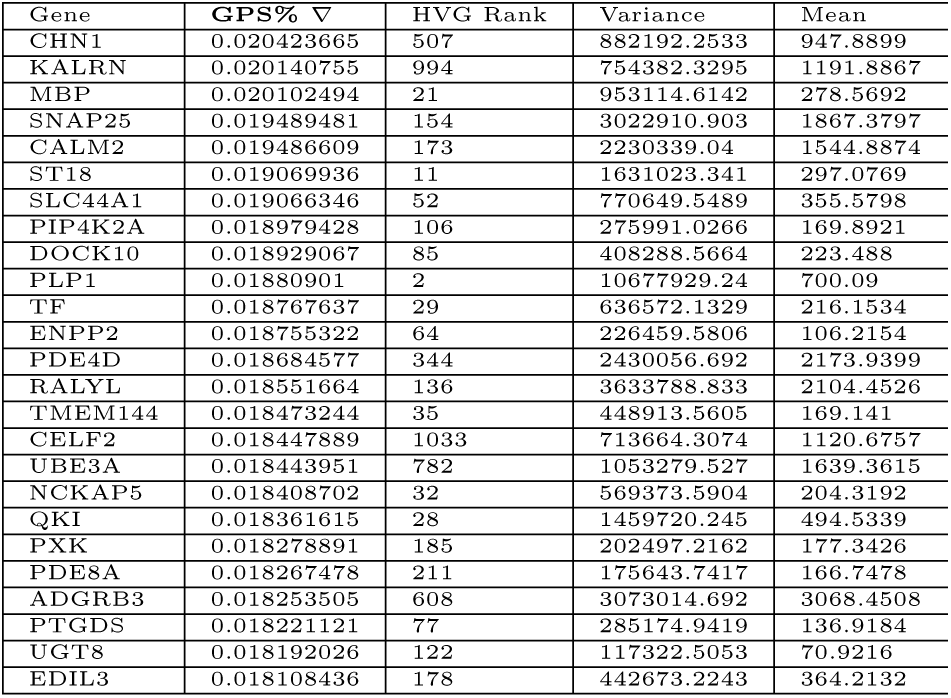
MMC Top 25 Principal Genes GPS and HVGs rank + Mean and Variance from count matrix.

**Table 3.**
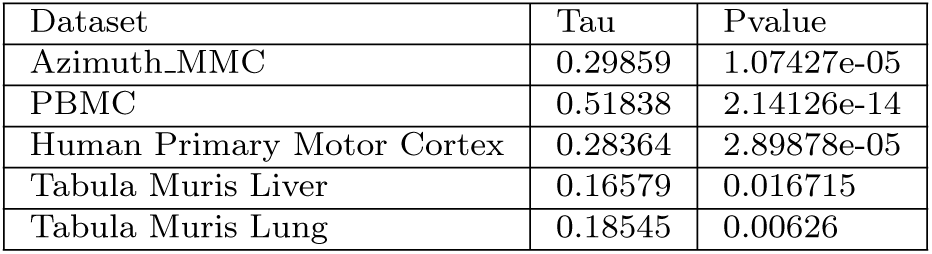
Kendall Rank Correlation Coefficient for the top 100 Principal Genes and Highly Variable Genes.

#### Biological characteristics in MMC data

As a showcase of utilizing PG in clustering, visualization and deriving biological insights from scRNA-Seq analysis, we show in Figure 10 the clustering results of MMC dataset using only top 5% of cumulative principal score with the resulting clusters composition match to the original cell types, with average accuracy of 91.69% and peak accuracy (0.965) observed at 2889 genes corresponding to 30% cumulative score. Indeed, the top 100 genes shown in the heatmap represent ‘important’ genes contributing to well separated cell partitioning. The enrichment analysis using only the top 100 PGs is shown in Table 4, highlighting the relevance of the selected genes for the mouse motor cortex neuronal cell types. Furthermore, the enrichment analysis, highlighting the biological characteristics and importance of the top selected genes for all the remaining datasets are given in section 0.5 of the supplementary materials.

**Figure 10.**
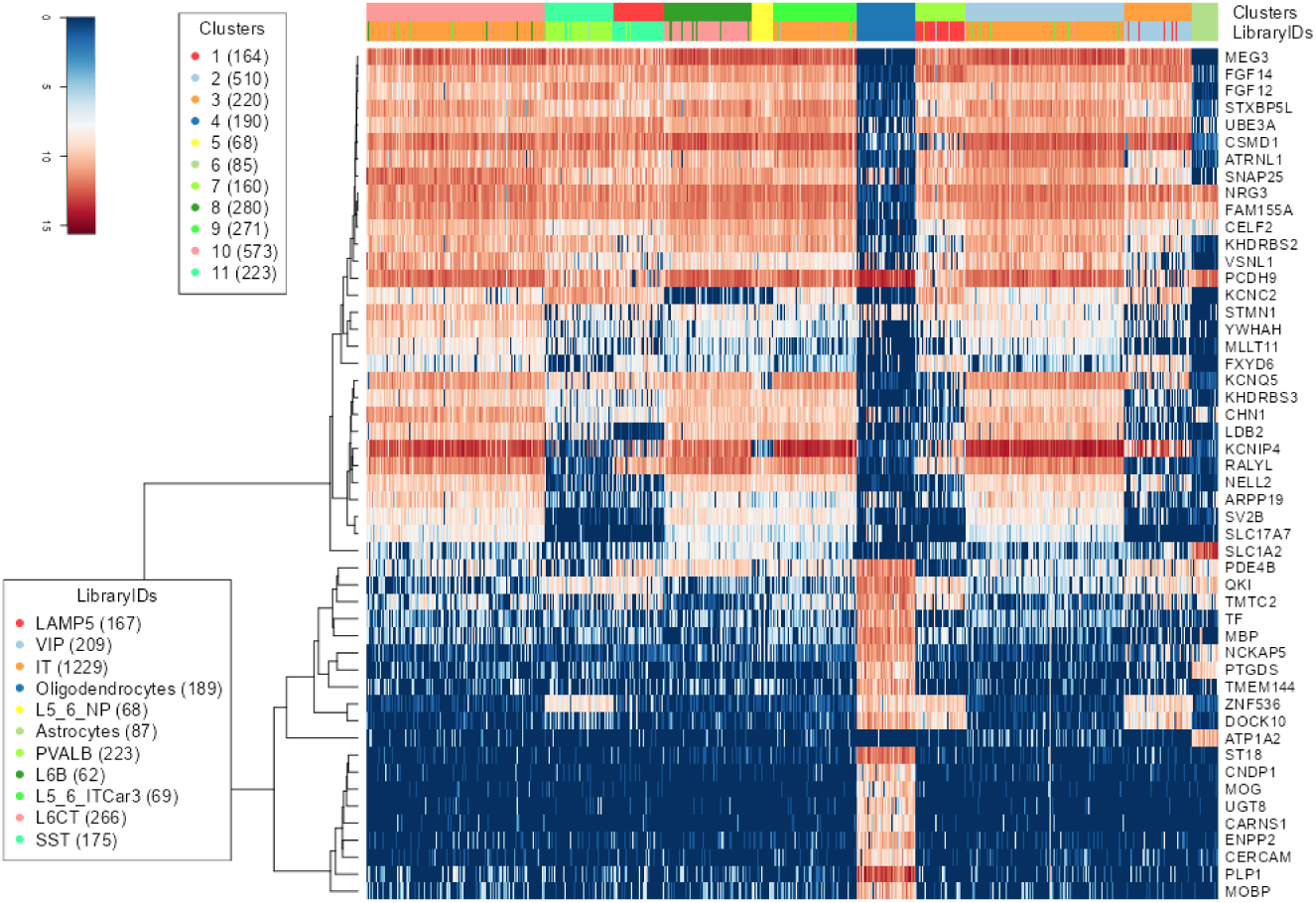
Heatmap representing the top 100 principal genes for MMC data showing the true and clustering labels. The selected PG genes clearly represent a set of genes with specificity to various cell types.

**Table 4.** Top 100 Principal Genes Enrichment Analysis for MMC dataset.

| Term ID | Term | NegLog10p | Intersection |
| --- | --- | --- | --- |
| GO:0007399 | nervous system development | 7.4558 | PLP1 DOCK10 KCNC2 LDB2 SNAP25 SLC1A2 ZNF536 MBP NRG3 QKI NELL2 CHN1 ENPP2 UBE3A STMN1 MOBP PCDH9 ATP1A2 UGT8 |
| GO:0048731 | system development | 5.7685 | MEG3 PLP1 DOCK10 KCNC2 LDB2 SNAP25 SLC1A2 ZNF536 MBP NRG3 QKI NELL2 CHN1 ENPP2 CSMD1 UBE3A STMN1 MOBP PCDH9 ATP1A2 UGT8 |
| GO:0046873 | metal ion transmembrane transporter activity | 5.6186 | KCNIP4 KCNQ5 KCNC2 SNAP25 SLC1A2 PDE4B ATP1A2 FGF12 FXYD6 YWHAH SLC17A7 |
| GO:0045202 | synapse | 5.4482 | DOCK10 KCNQ5 KCNC2 SNAP25 SLC1A2 NRG3 QKI PDE4B UBE3A PCDH9 ATP1A2 FGF12 STXBP5L FXYD6 SV2B YWHAH |
| GO:0008076 | voltage-gated potassium channel complex | 5.2160 | KCNIP4 KCNQ5 KCNC2 SNAP25 |
| GO:0034705 | potassium channel complex | 4.9541 | KCNIP4 KCNQ5 KCNC2 SNAP25 |
| GO:0022890 | inorganic cation transmembrane transporter activity | 4.6706 | KCNIP4 KCNQ5 KCNC2 SNAP25 SLC1A2 PDE4B ATP1A2 FGF12 FXYD6 YWHAH SLC17A7 |
| GO:0043005 | neuron projection | 4.5567 | KCNIP4 DOCK10 KCNQ5 KCNC2 SNAP25 SLC1A2 MBP NELL2 PDE4B STMN1 PCDH9 ATP1A2 |
| GO:0007275 | multicellular organism development | 4.4204 | MEG3 PLP1 DOCK10 KCNC2 LDB2 SNAP25 SLC1A2 ZNF536 MBP NRG3 QKI NELL2 CHN1 ENPP2 CSMD1 UBE3A STMN1 MOBP PCDH9 ATP1A2 UGT8 |
| GO:0008324 | monoatomic cation transmembrane transporter activity | 4.4114 | KCNIP4 KCNQ5 KCNC2 SNAP25 SLC1A2 PDE4B ATP1A2 FGF12 FXYD6 YWHAH SLC17A7 |

From the results shown throughout this section, PGs offer an interpretable approach to selecting highly variable genes without an arbitrary configuration for the number of top genes. The ranks of PGss show clear relevance to the variance and mean of the gene expression, a property not clearly shown for the top HVGs. Furthermore, higher accuracies can be achieved with a lower number of selected PGs, which allows for more reduction in the size of the high dimensional datasets and for more efficient downstream analyses.

### 3.5 Method performance

Figure 11 a) shows that the IRLBA algorithm vastly outperforms SVD for the calculation of the first few principal components with the highest associated eigenvalues. Both methods show a slight increase in computation time when increasing the number of principal components; typically though, a number less than 20 PCs can be used in our analysis as discussed in section 3.2.

**Figure 11.**
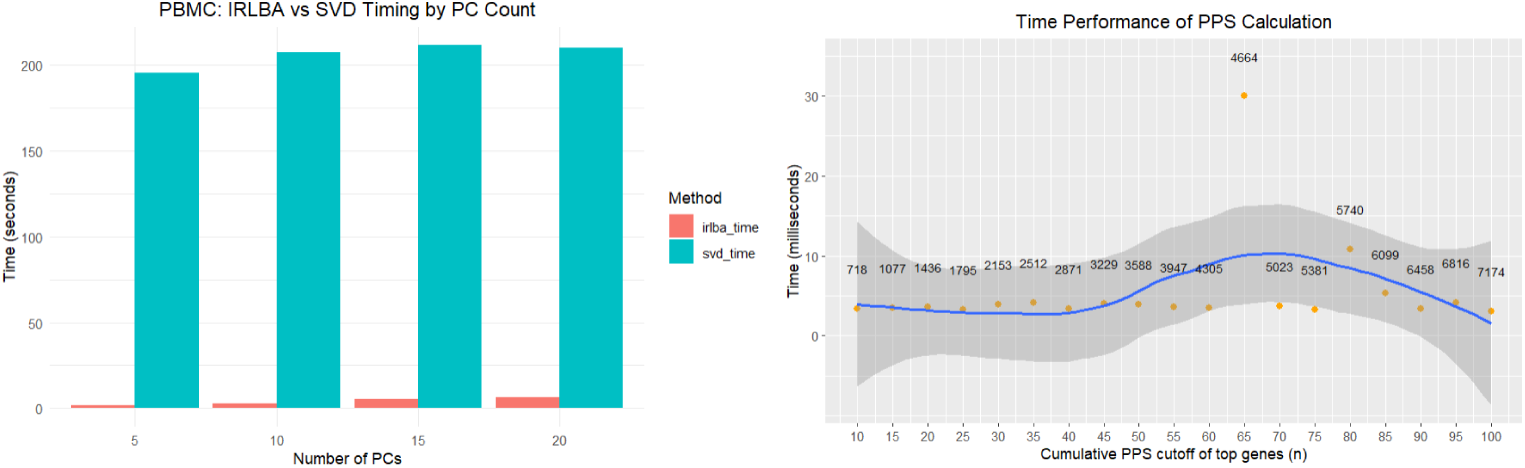
a) PCA performance time comparison for PBMCs (7174 genes *×* 2882 cells), IRLBA vs. SVD PCA algorithms in red and blue respectively. b) Time to calculate GPS for varied cumulative score cutoffs for PBMCs.

We also examined the additional time component needed for the calculation of GPS scores and gene selection based on the cumulative scores cutoff once the PCA step is concluded. Figure 11 b) shows the results for PBMC dataset. The additional time in this step is trivial in comparison to running the IRLBA or SVD algorithm and varies between 3 and 30 milliseconds. Interestingly, a drop in computation time is observed when selecting a high GPS cutoff, especially the full 100% since at that point the full list of genes is directly used.

## 4 Conclusion

. Our study introduced a novel method for selecting highly variable genes in scRNA-Seq datasets called *Principal Genes* that utilizes the rotations/loadings matrix data of PCA. We created a new scoring metric to quantify and rank Principal Genes variability called GPS score and used this approach to select the top Principal genes for further downstream analysis tasks. We validated our approach on multiple real scRNA-Seq datasets and showed that Principal genes are on par and often exceed the performance of dispersion-based HVG method. Our results show that using principal genes produces consistently high sensitivity and ARI scores for all tested datasets. Furthermore, our method is highly interpretable and flexible in adjusting the total % cumulative variability contribution cutoff, which in turn decides on the number of top Principal Genes without arbitrarily setting a number of genes to use. Finally, the method’s performance in terms of execution time is ultra-fast, especially for the GPS based Principal Genes selection step. Our method is a valuable addition to the repertoire of tools available for analyzing scRNA-Seq data and can easily be used in combination with other methods to achieve best possible results.

## Author Contributions

E.K.: Conceptualization, Software Implementation, Validation, Formal Analysis, Data Curation, Results Analysis, Writing-Original Draft, Writing- Review & Editing, Visualization. S.K.: Investigation, Data Curation T.N.: Code Review, Data Curation M.M.: Conceptualization, Methodology, Validation, Formal Analysis, Results Analysis, Writing-Original Draft, Writing- Review & Editing, Supervision, Project Administration, Funding Acquisition.

## Data Availability

Principal Genes algorithm used in this paper is available as an R project via GitHub repository at https://github.com/moussa-lab/PrincipalGenes and from the authors upon reasonable request.

## Acknowledgments

This work was supported by the National Science Foundation [NSF-2341725, NSF-2443386]; National Institutes of Health [NIH-K25CA270079]; and the University of Oklahoma Big Idea Challenge 2.0 award (BIC2.0). Research reported in this publication was also supported in part by the National Institute of General Medical Sciences of the National Institutes of Health under Award Number P20GM162339.

## Supplementary Materials

The following supporting information can be downloaded at the paper’s supplementary materials file: Figure S1: Bootstrapping results. Figures S2 and S3: VST Comparison. Figures S4 and S5: Results without housekeeping gene cleanup. Figures S6 to S10 Violin Plots for all datasets; Tables S1 to S5: Gene Enrichment Analyzes for all datasets. Table S6 and S7: Top 25 Shared Selected Genes in Azimuth’s MMC data (Intersect of Top 100 Principal Genes & Top 100 HVGs) and PBMCs

