## Supplemental Material for "Principal Genes: A PCA-based approach to highly variable genes selection for scRNA-Seq analysis"

### 0.1 Bootstrapping Results

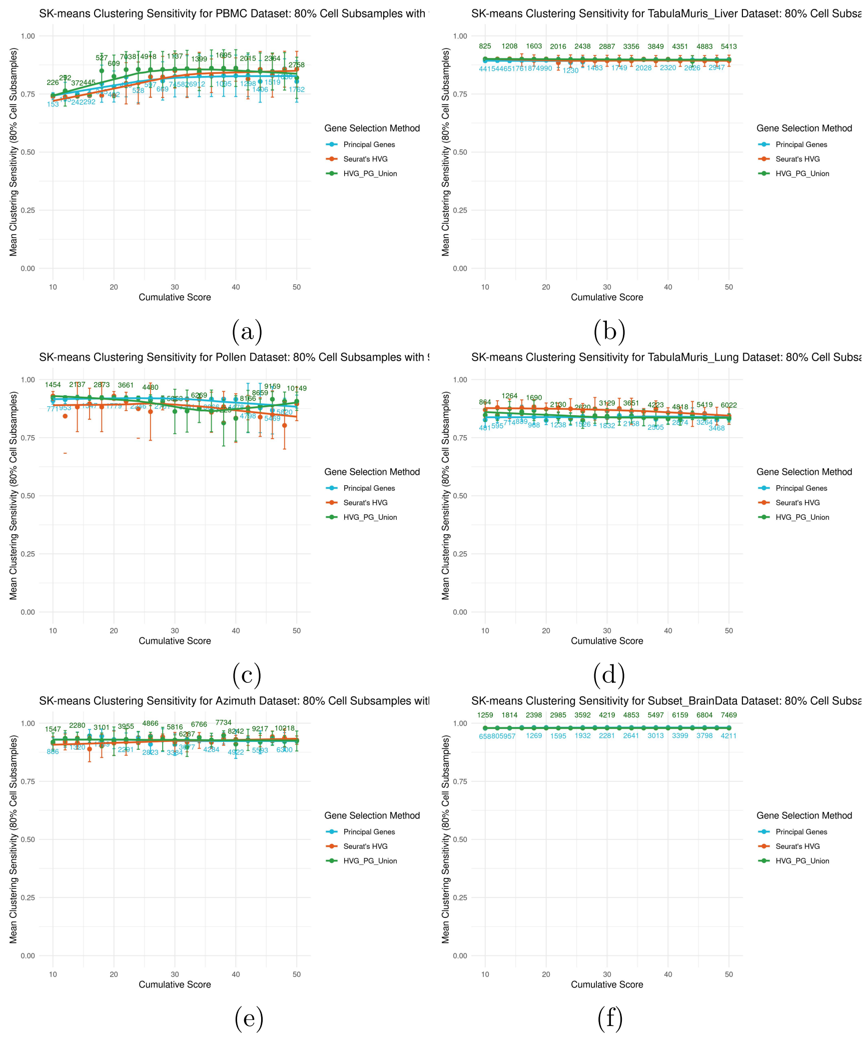


**Figure S1.** SK-Means Clustering Mean Sensitivity comparison between Principal Genes, Highly Variable Genes, and the union of both with 5% error bars for a) PBMCs, b) Tabula Muris Liver, c) Pollen, d) Tabula Muris Lung, e) Azimuth MMC and (f) Human Primary Motor Cortex. PGs selected for clustering are ordered in descending order and subset into groups of increasing cumulative GPS . HVGs are selected to match this number of genes in each group. The union is the sum of unique genes selected by both methods at each point. The number of genes at each group are shown in blue for HVGs and PGs and in green for the union.

### 0.2 Comparison with VST

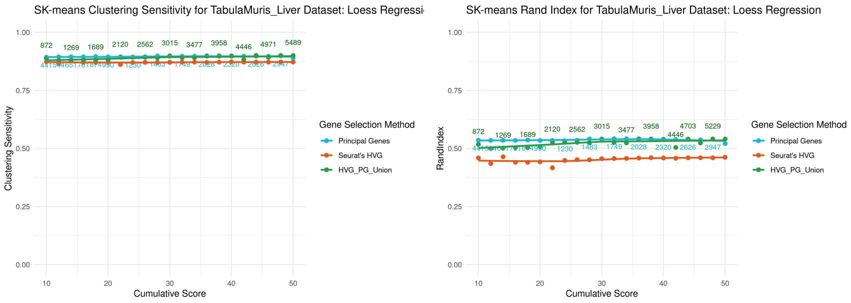


(a) (b)

**Figure S2.** Comparison between performance of Principal Genes and Highly Variable Genes selected via variance stabilization instead of disperion for Tabula Muris Liver dataset: a) SK-Means Clustering Sensitivity and b) Adjusted Rand index. PGs selected for clustering are ordered in descending order and subset into groups of increasing cumulative GPS . HVGs are selected to match this number of genes in each group. The union is the sum of unique genes selected by both methods at each point. The number of genes at each group are shown in blue for HVGs and PGs and in green for the union.

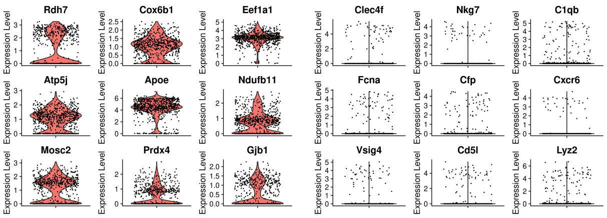


(a) (b)

**Figure S3.** Tabula Muris Liver Dataset run with variance stabilization instead of dispersion for Highy Variable Gene selection: a) Violin Plots for top 9 genes in PG's top 100 absent in HVG's top 100. b) Violin Plots for top 9 genes in HVG's top 100 absent in PG's top 100. No overlap in the top 100 PGs and top 100 HVGs.

### 0.3 Performance results without Housekeeping Gene Filtering

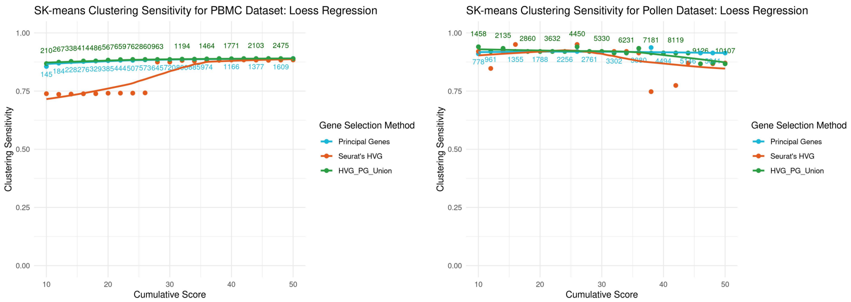


(a) (b)

**Figure S4.** SK-Means Clustering SessAdjusted Rand Index comparison between Principal Genes selected without filtering for housekeeping genes, Highly Variable Genes, and the union of both: a) PBMCs and b) Pollen. PGs selected for clustering are ordered in descending order and subset into groups of increasing cumulative GPS . HVGs are selected to match this number of genes in each group. The union is the sum of unique genes selected by both methods at each point. The number of genes at each group are shown in blue for HVGs and PGs and in green for the union.

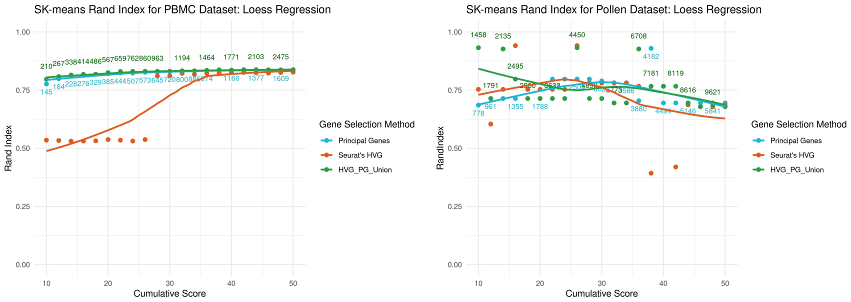


(a) (b)

**Figure S5.** Adjusted Rand Index comparison between Principal Genes selected without filtering for housekeeping genes, Highly Variable Genes, and the union of both: a) PBMCs and b) Pollen. PGs selected for clustering are ordered in descending order and subset into groups of increasing cumulative GPS . HVGs are selected to match this number of genes in each group. The union is the sum of unique genes selected by both methods at each point. The number of genes at each group are shown in blue for HVGs and PGs and in green for the union.

### 0.4 All datasets' violin plots

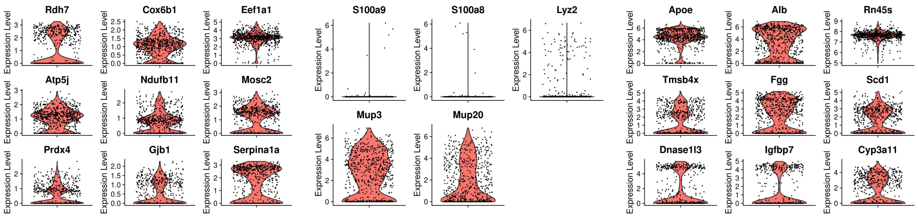


(a) (b) (c)

**Figure S6.** Tabula Muris Liver Dataset: a) Violin Plots for top 9 genes in PG's top 100 absent in HVG's top 100. b) Violin Plots for top 5 genes in HVG's top 100 absent in PG's top 100. c) Violin Plots for 9 genes in the intersection of HVG's and PG's top 100.

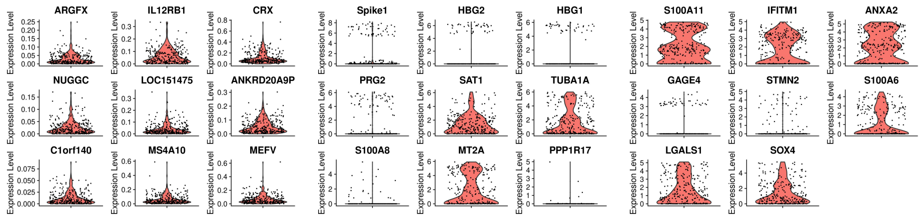


(a) (b) (c)

**Figure S7.** Pollen Dataset: a) Violin Plots for top 9 genes in PG's top 100 absent in HVG's top 100. b) Violin Plots for top 9 genes in HVG's top 100 absent in PG's top 100. c) Violin Plots for 8 genes in the intersection of HVG's and PG's top 100.

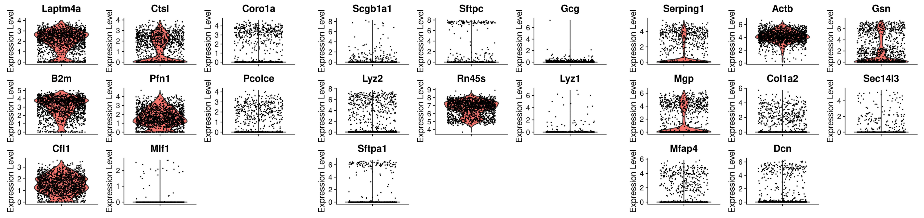


(a) (b) (c)

**Figure S8.** Tabula Muris Lung Dataset: a) Violin Plots for top 8 genes in PG's top 100 absent in HVG's top 100. b) Violin Plots for top 7 genes in HVG's top 100 absent in PG's top 100. c) Violin Plots for 8 genes in the intersection of HVG's and PG's top 100.

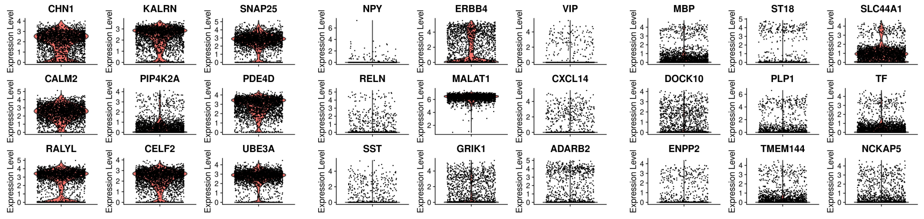


(a) (b) (c)

**Figure S9.** Azimuth MMC Dataset: a) Violin Plots for top 9 genes in PG's top 100 absent in HVG's top 100. b) Violin Plots for top 9 genes in HVG's top 100 absent in PG's top 100. c) Violin Plots for 9 genes in the intersection of HVG's and PG's top 100.

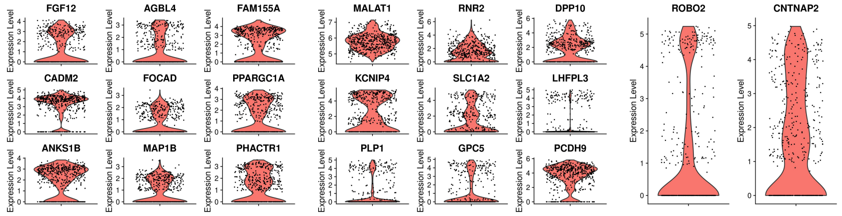


(a) (b) (c)

**Figure S10.** Human Primary Motor Cortex Dataset: a) Violin Plots for top 9 genes in PG's top 100 absent in HVG's top 100. b) Violin Plots for top 9 genes in HVG's top 100 absent in PG's top 100. c) Violin Plots for 2 genes in the intersection of HVG's and PG's top 100.

### 0.5 All datasets' Gene Enrichment Tables

**Table S1.** Top 100 Principal Genes Enrichment Analysis for PBMC dataset.

| Term ID | Term | NegLog10p | Intersection |
|---|---|---|---|
| GO:0006955 | immune response | 21.4007 | CD74 TYROBP GNLY CD79A CD79B S100A9 FCER1G CST3 GZMB AIF1 FTH1 LYZ NKG7 FCN1 CD3D MS4A1 CFP S100A8 JUNB LTB LST1 CD3E CST7 GZMA IL32 CYBA HCST CFD CTSW CD8B CD14 LY86 GRN SPI1 GPX1 BANK1 PRF1 FCGRT CD27 CD7 KLRB1 KLRF1 ACTG1 SPON2 FCGR3A CCL4 CD247 |
| GO:0140507 | granzyme-mediated programmed cell death signaling pathway | 6.6767 | GZMB NKG7 GZMA PRF1 SRGN |
| GO:0005515 | protein binding | 4.1396 | CD74 TYROBP GNLY CD79A CD79B S100A9 EEF1A1 FCER1G CST3 GZMB AIF1 FTH1 LYZ NKG7 FCN1 CD3D CD37 MS4A1 CFP MS4A6A S100A8 LDHB JUNB LTB HOPX CD3E COTL1 PFN1 TMEM176B CST7 TYMP GSTP1 GZMA IL32 CYBA HCST CD8B NOSIP CD14 LY86 GRN ACTB SPI1 SERPINA1 FGFBP2 GPX1 BANK1 EEF1D CFL1 TPT1 PRF1 CLIC3 CSF3R FCGRT CD27 IFI30 EEF1B2 PABPC1 NPM1 S100A4 CD7 KLRB1 TMEM176A SRGN KLRF1 LGALS2 S100A6 CEBPD ACTG1 NACA COX4I1 CLIC1 MYL12A SPON2 FCGR3A SPIB OAZ1 CCL4 CD247 |
| GO:0019865 | immunoglobulin binding | 2.8173 | FCER1G MS4A1 FCGRT FCGR3A |
| GO:0070661 | leukocyte proliferation | 2.7234 | CD74 TYROBP CD79A AIF1 MS4A1 JUNB LST1 CD3E GSTP1 FCGR3A |
| GO:0050786 | RAGE receptor binding | 2.4131 | S100A9 S100A8 S100A4 |
| GO:0048144 | fibroblast proliferation | 2.0784 | CD74 FTH1 GSTP1 GPX1 IFI30 S100A6 |
| GO:0006909 | phagocytosis | 2.0269 | TYROBP FCER1G AIF1 FCN1 CFP CYBA CD14 SPON2 |
| GO:0071621 | granulocyte chemotaxis | 1.7908 | CD74 S100A9 FCER1G S100A8 CSF3R CCL4 |
| GO:0070488 | neutrophil aggregation | 1.5323 | S100A9 S100A8 |
| GO:0045730 | respiratory burst | 1.5301 | S100A9 CYBA GRN CD52 |
| GO:0003746 | translation elongation factor activity | 1.3201 | EEF1A1 EEF1D EEF1B2 |
| GO:0035662 | Toll-like receptor 4 binding | 1.3051 | S100A9 S100A8 |

**Table S2.** Top 100 Principal Genes Enrichment Analysis for Tabula Muris Liver dataset.

| Term ID | Term | NegLog10p | Intersection |
|---|---|---|---|
| GO:0045333 | cellular respiration | 7.1486 | COX6B1 NDUFB11 CISD1 UQCRFS1 NDUFA5 UQCR10 FBP1 NDUFA13 CHCHD10 SDHC COX7B TPI1 COX6A1 |
| GO:0044281 | small molecule metabolic process | 6.8510 | RDH7 APOE NDUFB11 C3 APOA5 CTH NDUFA5 GSTZ1 FBP1 SLC27A2 NDUFA13 SCP2 ACSL1 SEPHS2 GPX1 SCD1 ASS1 PON1 IGF1 SORD SDHC BOLA3 FDFT1 TPI1 |
| GO:0006958 | complement activation, classical pathway | 4.2272 | C3 C4BP CFI SERPING1 MBL2 |
| GO:0002526 | acute inflammatory response | 3.2341 | SERPINA1A C3 ITIH4 ASS1 DNASE1L3 FN1 |
| GO:0006641 | triglyceride metabolic process | 2.9472 | APOE C3 APOA5 ACSL1 GPX1 SCD1 |
| GO:0010876 | lipid localization | 2.8601 | APOE C3 APOA5 SLC27A2 SCP2 ACSL1 SLCO1B2 APOM PON1 PLIN2 |
| GO:0006952 | defense response | 2.7766 | APOE SERPINA1A C3 C4BP CFI FGA ITIH4 SERPING1 B2M FGG CD1D1 GPX1 SCD1 ASS1 MBL2 DNASE1L3 FN1 DDT FAU |
| GO:0051050 | positive regulation of transport | 2.1557 | APOE C3 PGRMC1 B2M FGG SCP2 ACSL1 MBL2 PON1 IGF1 CHCHD10 FXYD1 |
| GO:0050197 | phytanate-CoA ligase activity | 2.1291 | SLC27A2 ACSL1 |
| GO:0070251 | pristanate-CoA ligase activity | 2.1291 | SLC27A2 ACSL1 |
| GO:0044419 | biological process involved in interspecies interaction between organisms | 1.9457 | APOE C3 C4BP CFI FGA SERPING1 HSP90AB1 B2M FGG CD1D1 GPX1 SCD1 SLCO1B2 ASS1 MBL2 FN1 CYP3A11 FAU |
| GO:0061134 | peptidase regulator activity | 1.6915 | SERPINA1A C3 ITIH4 SERPING1 SERPIND1 ITIH2 FN1 |
| GO:0045428 | regulation of nitric oxide biosynthetic process | 1.5658 | APOE HSP90AB1 ASS1 IGF1 |
| GO:0070542 | response to fatty acid | 1.5291 | ACSL1 SCD1 ASS1 BOLA3 |
| GO:0061136 | regulation of proteasomal protein catabolic process | 1.5284 | APOE UBB GABARAP HSP90AB1 GPX1 HERPUD1 |
| GO:0004866 | endopeptidase inhibitor activity | 1.5042 | SERPINA1A C3 ITIH4 SERPING1 SERPIND1 ITIH2 |
| GO:0042060 | wound healing | 1.4159 | APOE FGA HABP2 SERPING1 FGG SERPIND1 GPX1 FN1 |
| GO:0015909 | long-chain fatty acid transport | 1.3570 | APOE SLC27A2 ACSL1 PLIN2 |
| GO:0140077 | positive regulation of lipoprotein transport | 1.3013 | APOE PGRMC1 |

**Table S3.** Top 100 Principal Genes Enrichment Analysis for Pollen dataset.

| Term ID | Term | NegLog10p | Intersection |
|---|---|---|---|
| GO:0005509 | calcium ion binding | 4.0858 | S100A11 MYOF ANXA5 PCDH11Y PLSCR1 MYL4 ANXA2 S100A6 FSTL1 CALU FBN1 MYL12A ITGB1 |
| GO:0009611 | response to wounding | 3.0129 | MYOF ANXA5 NFIB PTPRS PLSCR1 ANXA2 MAP1B TPM1 SCARF1 MYL12A ITGB1 |
| GO:0065007 | biological regulation | 2.7747 | ARGFX IL12RB1 S100A11 CRX NUGGC MS4A10 MEFV TMSB10 KCNJ5 MYOF PRAME IFITM1 ANXA5 NFIB FOXK1 DKK1 CRTAM PCDH11Y ADRA1A PTPRS HTRA4 FBLIM1 NLRP12 ALCAM ZNF713 BLVRB PLSCR1 MYL4 ANXA2 STMN2 MAP1B NYAP2 S100A6 FRMD6 FSTL1 NMUR1 TPM1 LGALS1 FBN1 ODF2L DAND5 MAGEA12 EPHA10 SCARF1 IFI6 SOX4 RNF144B KCNJ11 MYL12A ADAMTSL2 NEK7 ITGB1 LY6E |
| GO:0098609 | cell-cell adhesion | 2.3471 | IL12RB1 S100A11 CRTAM PCDH11Y PTPRS FBLIM1 ALCAM ANXA2 LGALS1 SCARF1 SOX4 MYL12A ITGB1 |
| GO:0044548 | S100 protein binding | 2.2936 | S100A11 ANXA2 S100A6 |
| GO:0050896 | response to stimulus | 2.1361 | IL12RB1 S100A11 NUGGC MS4A10 MEFV MYOF PRAME IFITM1 ANXA5 NFIB FOXK1 DKK1 CRTAM PCDH11Y ADRA1A PTPRS HTRA4 NLRP12 ALCAM BLVRB PLSCR1 ANXA2 STMN2 MAP1B NYAP2 S100A6 FRMD6 FSTL1 NMUR1 TPM1 LGALS1 FBN1 DAND5 EPHA10 SCARF1 IFI6 SOX4 KCNJ11 MYL12A ADAMTSL2 NEK7 ITGB1 LY6E |
| GO:0048468 | cell development | 2.0586 | IL12RB1 NFIB DKK1 CRTAM ADRA1A FUT6 PTPRS ALCAM BLVRB ANXA2 STMN2 MAP1B NYAP2 S100A6 FRMD6 TPM1 LGALS1 FBN1 EPHA10 SCARF1 SOX4 ITGB1 |
| GO:0008092 | cytoskeletal protein binding | 1.7625 | MEFV TMSB10 FBLIM1 MYL4 ANXA2 STMN2 MAP1B S100A6 TPM1 KCNJ11 MYL12A ITGB1 |
| GO:0007154 | cell communication | 1.7209 | IL12RB1 S100A11 MS4A10 MEFV KCNJ5 MYOF PRAME IFITM1 ANXA5 DKK1 PCDH11Y ADRA1A PTPRS HTRA4 NLRP12 ALCAM BLVRB PLSCR1 MAP1B NYAP2 S100A6 FRMD6 FSTL1 NMUR1 LGALS1 FBN1 DAND5 EPHA10 IFI6 SOX4 KCNJ11 ADAMTSL2 NEK7 ITGB1 LY6E |
| GO:0000902 | cell morphogenesis | 1.4306 | NFIB DKK1 PTPRS FBLIM1 ALCAM MAP1B NYAP2 S100A6 FRMD6 TPM1 EPHA10 ITGB1 |
| GO:0001996 | positive regulation of heart rate by epinephrine-norepinephrine | 1.3139 | ADRA1A TPM1 |

**Table S4.** Top 100 Principal Genes Enrichment Analysis for Tabula Muris Lung dataset.

| Term ID | Term | NegLog10p | Intersection |
|---|---|---|---|
| GO:0005201 | extracellular matrix structural constituent | 12.3439 | COL6A1 COL1A2 SPARC DCN LTBP4 BGN FBLN1 COL6A2 ECM1 DPT COL1A1 NID1 IGFBP6 |
| GO:0048870 | cell motility | 9.4311 | ACTB CORO1A CFL1 GNAI2 RAC1 TMSB4X SPARC DCN P4HB TUBB4B FCER1G FBLN1 FSTL1 CAV1 TEKT1 ACVRL1 NME5 ECM1 GPX1 COL1A1 MYADM TUBA1A S1PR1 FGFR1 NRP1 IGFBP6 CYGB PFN1 |
| GO:0065008 | regulation of biological quality | 8.7238 | SERPING1 CORO1A COL6A1 CFL1 GNAI2 COL1A2 RAC1 GSN ARPC2 DCN P4HB UBB LTBP4 FCER1G FBLN1 SCPEP1 APP CYBA CTSL CAV1 TY-ROBP LAMP1 CYP2S1 ACVRL1 RBP1 GPX1 MYADM GAPDH YBX1 TUBA1A CD47 FGFR1 NRP1 THBD PFN1 |
| GO:0005515 | protein binding | 8.2815 | ACTB CORO1A COL6A1 CFL1 B2M GNAI2 EEF1A1 COL1A2 TIMP2 RAC1 TMSB4X GSN SERPINH1 SPARC ARPC2 DCN ARPC1B PCOLCE P4HB UBB SPARCL1 LTBP4 BGN LAPTM5 FCER1G OLFML2B TPPP3 FBLN1 APP FANK1 CYBA CTSL FSTL1 CAV1 TYROBP LAMP1 LRRC10B ACVRL1 EHD2 PSAP ECM1 COTL1 HTRA3 UBXN10 GPX1 COL1A1 FAM81A GAPDH YBX1 TUBA1A MLF1 NID1 S1PR1 CD47 ARPC4 FGFR1 NRP1 SPAG17 H3F3B IGFBP6 CTSS ARHGDIG HMGCS2 PFN1 ZMYND10 ISLR |
| GO:0045229 | external encapsulating structure organization | 8.2258 | COL6A1 COL1A2 SERPINH1 LTBP4 OLFML2B FBLN1 ADAMTS2 APP CAV1 DPT COL1A1 NID1 CTSS |
| GO:0032963 | collagen metabolic process | 7.7760 | COL6A1 RCN3 COL1A2 SERPINH1 ADAMTS2 CTSL COL1A1 CTSS CYGB |
| GO:0012501 | programmed cell death | 3.4770 | ACTB CORO1A COL6A1 GNAI2 GSN P4HB UBB FCER1G APP CTSL FSTL1 CAV1 LAMP1 COL6A2 NME5 GPX1 GAPDH TUBA1A CD47 FGFR1 NRP1 PMP22 |
| GO:0045010 | actin nucleation | 3.0327 | CORO1A GSN ARPC2 ARPC1B ARPC4 |
| GO:0005200 | structural constituent of cytoskeleton | 2.5461 | ACTB ARPC2 TUBB4B TUBA1A ARPC4 |
| GO:0005539 | glycosaminoglycan binding | 2.5106 | DCN PCOLCE BGN APP FSTL1 FGFR1 NRP1 |
| GO:0005509 | calcium ion binding | 2.4487 | RCN3 GSN SPARC SPARCL1 LTBP4 FBLN1 FSTL1 EHD2 C1RA CALU NID1 THBD |
| GO:0031579 | membrane raft organization | 2.4023 | COL6A1 GSN CAV1 MYADM |
| GO:0051604 | protein maturation | 1.7913 | B2M GSN SERPINH1 P4HB FBLN1 ADAMTS2 CTSL CTSS ZMYND10 |
| GO:0061134 | peptidase regulator activity | 1.6915 | SERPING1 TIMP2 SERPINH1 PCOLCE FBLN1 APP CAV1 |
| GO:0042110 | T cell activation | 1.6406 | ACTB CORO1A B2M GSN LAPTM5 FCER1G CTSL CAV1 TYROBP PSAP |
| GO:0007166 | cell surface receptor signaling pathway | 1.5311 | CORO1A COL6A1 GNAI2 COL1A2 RAC1 DCN LTBP4 LAPTM5 FCER1G APP CYBA CAV1 TYROBP ACVRL1 ECM1 HTRA3 GPX1 COL1A1 TUBA1A NID1 FGFR1 NRP1 |
| GO:0070887 | cellular response to chemical stimulus | 1.5094 | ACTB CORO1A COL6A1 B2M GNAI2 COL1A2 RAC1 P4HB FCER1G CAV1 PSAP GPX1 COL1A1 TUBA1A S1PR1 FGFR1 NRP1 CTSS HMGCS2 CYGB |
| GO:0019882 | antigen processing and presentation | 1.4898 | B2M FCER1G CTSL PSAP CTSS |
| GO:0000302 | response to reactive oxygen species | 1.4395 | COL6A1 NME5 PSAP GPX1 COL1A1 CYGB |
| GO:0021872 | forebrain generation of neurons | 1.4039 | B2M RAC1 UBB FGFR1 NRP1 |

**Table S5.** Top 100 Principal Genes Enrichment Analysis for Human Primary Motor Cortex dataset.

| Term ID | Term | NegLog10p | Intersection |
|---|---|---|---|
| GO:0048666 | neuron development | 11.1208 | ROBO2 AGBL4 MAP1B PHACTR1 CHN1 MAGI2 APP MYT1L FAT3 IQSEC1 NELL2 CNTNAP2 FRY OLFM3 SPTBN4 CNTN1 NRP1 CAMK2A CSMD3 NEDD4L NTNG1 ARHGAP44 SLIT3 EPB41L3 STAU2 PAK1 AUTS2 DAB1 |
| GO:0008093 | cytoskeletal anchor activity | 2.9406 | BICD1 ANK2 SPTBN4 EPB41L3 |
| GO:0051049 | regulation of transport | 2.9240 | FGF12 MAP1B GRIN2A MAGI2 BICD1 ANK2 PRKAG2 SPTBN4 CNTN1 NRP1 SNAP91 CAMK2A ARHGEF7 ITPR1 CACNA1D NEDD4L GRIA1 ARHGAP44 MKLN1 SYBU CDH13 |
| GO:0030029 | actin filament-based process | 2.8542 | FGF12 PHACTR1 IQSEC1 ANK2 SPTBN4 NRP1 CACNA1D NEDD4L ARHGAP44 EPB41L3 MKLN1 STAU2 PAK1 AUTS2 TTC17 |
| GO:0008092 | cytoskeletal protein binding | 2.8405 | AGBL4 MAP1B PHACTR1 BICD1 SYNE1 STRBP ANK2 EML6 SPTBN4 ARHGEF7 TTLL11 CACNA1D GRIA1 EPB41L3 STAU2 PAK1 |
| GO:1990806 | ligand-gated ion channel signaling pathway | 2.7890 | GRIN2A APP ITPR1 GRIA1 GRID1 |
| GO:0010646 | regulation of cell communication | 2.4630 | ROBO2 FGF12 MAP1B CHN1 GRIN2A MAGI2 APP BICD1 IQSEC1 RGS7 ANK2 NRP1 CAMK2A TSC22D1 ARHGEF7 ITPR1 NTNG1 GRIA1 ARHGAP44 SLIT3 SHANK2 PPP1R16B GRID1 STAU2 PAK1 NLK AUTS2 DYNC2H1 SYBU CDH13 DAB1 TNKS |
| GO:0019899 | enzyme binding | 1.9114 | PPARGC1A MAGI2 APP BICD1 PEX5L SYNE1 ANK2 NELL2 CNTNAP2 PRKAG2 SPTBN4 NRP1 SNAP91 RAPGEF6 ARHGEF7 GRIA1 ARHGAP44 CHCHD3 RABGAP1L STAU2 PAK1 NLK |
| GO:0010970 | transport along microtubule | 1.8812 | AGBL4 MAP1B APP BICD1 STAU2 DYNC2H1 SYBU |
| GO:0004970 | glutamate-gated receptor activity | 1.5726 | GRIN2A GRIA1 GRID1 |
| GO:0099536 | synaptic signaling | 1.4890 | FGF12 MAP1B GRIN2A APP CAMK2A ITPR1 NTNG1 GRIA1 ARHGAP44 SHANK2 GRID1 STAU2 SYBU |

### 0.6 Top 25 Shared Genes Ranking

**Table S6.** Top 25 Shared Selected Genes in PBMCs data: Intersect of Top 100 Principal Genes & Top 100 HVGs

| Gene | GPS_Rank | Variance% ∇ | Mean | HVG_Rank |
|---|---|---|---|---|
| GNLY | 6 | 173.1488 | 8.2321 | 1 |
| CD74 | 2 | 80.2271 | 3.8536 | 2 |
| NKG7 | 21 | 69.157 | 5.3567 | 9 |
| ACTB | 56 | 39.4282 | 10.7384 | 49 |
| HLA_DRA | 1 | 21.8914 | 1.628 | 8 |
| HLA_DRB1 | 8 | 13.8246 | 1.3584 | 11 |
| CD52 | 55 | 8.6412 | 3.161 | 76 |
| TYROBP | 3 | 8.5391 | 1.7734 | 23 |
| GZMB | 17 | 7.2805 | 1.2866 | 21 |
| S100A9 | 10 | 6.9545 | 0.4289 | 3 |
| CST3 | 16 | 5.6507 | 0.4285 | 5 |
| HCST | 47 | 5.6315 | 2.0885 | 86 |
| GZMA | 43 | 5.4048 | 1.3383 | 30 |
| FCER1G | 15 | 4.8345 | 1.2391 | 37 |
| CD37 | 24 | 4.2072 | 1.6013 | 74 |
| S100A8 | 29 | 3.378 | 0.2856 | 7 |
| CD79A | 7 | 3.3343 | 0.5385 | 17 |
| CD3D | 23 | 2.9197 | 1.3074 | 98 |
| HOPX | 35 | 2.3595 | 0.8043 | 60 |
| AIF1 | 18 | 1.7441 | 0.356 | 20 |
| FCN1 | 22 | 0.8435 | 0.1523 | 15 |
| HLA_DQA2 | 14 | 0.7372 | 0.2582 | 65 |
| TYMP | 41 | 0.4401 | 0.1613 | 54 |
| CFP | 27 | 0.2126 | 0.076 | 61 |
| SPI1 | 57 | 0.1553 | 0.0715 | 100 |

**Table S7.** Shared Selected Genes in Azimuth's MMC data: Intersect of Top 100 Principal Genes & Top 100 HVGs

| Gene | GPS_Rank | Variance% ∇ | Mean | HVG_Rank |
|---|---|---|---|---|
| KCNIP4 | 34 | 39404306.84 | 6402.2001 | 13 |
| PCDH9 | 69 | 11516891.43 | 4269.3637 | 42 |
| PLP1 | 10 | 10677929.24 | 700.09 | 2 |
| IL1RAPL1 | 38 | 4057983.973 | 1942.5295 | 70 |
| SLC1A2 | 63 | 3037474.873 | 355.6097 | 3 |
| ST18 | 6 | 1631023.341 | 297.0769 | 11 |
| QKI | 19 | 1459720.245 | 494.5339 | 28 |
| PDE4B | 48 | 1256151.116 | 586.5277 | 62 |
| MBP | 3 | 953114.6142 | 278.5692 | 21 |
| SLC44A1 | 7 | 770649.5489 | 355.5798 | 52 |
| TMTC2 | 43 | 739903.7797 | 353.3881 | 60 |
| RNF220 | 42 | 676266.1262 | 204.7843 | 23 |
| CTNNA3 | 39 | 670383.0206 | 258.9176 | 43 |
| TF | 11 | 636572.1329 | 216.1534 | 29 |
| NCKAP5 | 18 | 569373.5904 | 204.3192 | 32 |
| TMEM144 | 15 | 448913.5605 | 169.141 | 35 |
| ADGRV1 | 30 | 421349.2234 | 138.5066 | 24 |
| DOCK10 | 9 | 408288.5664 | 223.488 | 85 |
| PTGDS | 23 | 285174.9419 | 136.9184 | 77 |
| MOBP | 77 | 270328.3645 | 116.105 | 56 |
| ENPP2 | 12 | 226459.5806 | 106.2154 | 64 |
| CNDP1 | 64 | 164958.352 | 81.3309 | 59 |
| ATP1A2 | 92 | 104692.3031 | 51.422 | 54 |
